# Mini-batch optimization enables training of ODE models on large-scale datasets

**DOI:** 10.1101/859884

**Authors:** Paul Stapor, Leonard Schmiester, Christoph Wierling, Bodo M.H. Lange, Daniel Weindl, Jan Hasenauer

## Abstract

Quantitative dynamical models are widely used to study cellular signal processing. A critical step in modeling is the estimation of unknown model parameters from experimental data. As model sizes and datasets are steadily growing, established parameter optimization approaches for mechanistic models become computationally extremely challenging. However, mini-batch optimization methods, as employed in deep learning, have better scaling properties. In this work, we adapt, apply, and benchmark mini-batch optimization for ordinary differential equation (ODE) models thereby establishing a direct link between dynamic modeling and machine learning. On our main application example, a large-scale model of cancer signaling, we benchmark mini-batch optimization against established methods, achieving better optimization results and reducing computation by more than an order of magnitude. We expect that our work will serve as a first step towards mini-batch optimization tailored to ODE models and enable modeling of even larger and more complex systems than what is currently possible.

## Introduction

Cellular signal processing controls key properties of diverse mechanisms such as cell division (42), growth (36), differentiation (7), or apoptosis (56). Understanding its highly dynamic and complex nature is one of the major goals of systems biology (29). A common approach is modeling signaling pathways using ordinary differential equations (ODEs) (10; 14; 27; 60; 68). To account for the complex cross-talk between different pathways, recent models grew increasingly large, reaching the boundaries of what is currently computationally feasible (14; 21; 31; 54).

Most ODE models contain unknown parameters, e.g., reaction rate constants, which have to be inferred from measurement data such as immuno-blotting (8), proteomics (21), quantitative PCR (3), or viability (14). The larger a model becomes, the more data are needed to ensure the reliability of parameter estimates and model predictions (2). For models of cancer signaling, public databases (4; 12; 35; 43) can be exploited. Yet, if the data used to calibrate the model is derived from different experimental conditions, those constitute independent initial value problems, which need to be simulated repeatedly during model training (48). This causes a linear scaling of the computation time with the number of experimental conditions to be simulated. For large-scale ODE models with hundreds to thousands of chemical species and thousands of experimental conditions, this can take tens of thousands of computing hours, even if one applies state-of-the-art methods, such as exploiting adjoint sensitivity analysis and hierarchical problem structures (14; 54).

For training of ODE models, gradient-based approaches such as multi-start local optimization (48) or hybrid scatter search (62) have repeatedly shown to be the best performing methods to date (48; 52; 62). In multi-start local optimization, local optimizations are initialized at many randomized starting points in order to globally explore the parameter space (15). For small- to medium-scale models, these methods unravel the structure of local optima and recover the same global optimum (20; 48) reproducibly. However, for large-scale models, where each local optimization is computationally expensive, only a small number of starts can be performed (14; 21; 54). This is one of the main reasons why satisfactory parameter optimization for large-scale ODE models is still an open problem (26).

In the field of deep learning, where also gradient-based local optimization methods are used (33; 40; 51; 58), model training is often performed on vast datasets, requiring many independent model evaluations (25; 32). The problematic scaling behavior with respect to the size of the dataset is addressed by mini-batch optimization (1; 19; 57; 66): In each step of parameter optimization, a randomly sampled subset – a mini-batch – of the training data is used to inform the optimization process (19; 49). Hence, the model is only evaluated on a fraction of the dataset per optimization step, which leads to a drastic reduction of computation time (19; 66).

Sophisticated implementations of many mini-batch optimization algorithms are available in state-of-the-art toolboxes for neural nets, such as Tensorflow (1). Conceptually, these frameworks can be employed to mimic simple ODE solver schemes, e.g., a forward Euler integration, such as done in (67). However, it is well-known that ODE models in systems biology typically exhibit stiff dynamics. This makes it necessary to employ advanced numerical integration methods, such as implicit solvers with adaptive time stepping (30; 41; 48). This implies that it is essential to combine advanced methods from both fields, deep learning and ODE modeling. Furthermore, it is not clear how hyperparameters of mini-batch optimization methods, such as the mini-batch size, the learning rate, the optimization algorithm or other tuning parameters affect the optimization process for ODE models.

We implemented various mini-batch optimization algorithms for ODE models. We benchmarked these algorithms on small- to medium-scale ODE models, identified the most important hyperparameters for successful parameter optimization, and introduced algorithmic improvements, which are tailored to ODE modeling. Then, we transfer the approach to a large-scale model of cancer signaling (14), which we trained on a dataset comprising 13,000 experimental conditions – an unprecedented scale for training an ODE model. For this application example, we benchmarked our approach against state-of-the-art methods (14), achieving better optimization results while reducing the computation time by more than an order of magnitude. To the best of our knowledge, this is the first study integrating advanced training algorithms from deep learning with sophisticated tools from ODE modeling.

## Results

### Implementation of mini-batch optimization algorithms for ODE models

The inference of model parameters *θ* from experimental data is based on reducing a distance measure between simulated model outputs and measurements. In practice, the distance metric is often based on the assumption that the measurement noise is normally distributed and independent for each data point. The corresponding negative log-likelihood function *J*, which serves as objective or cost function, is (up to a constant, more details in the Methods section and the Supplementary Information) given by the sum of weighted least squares:

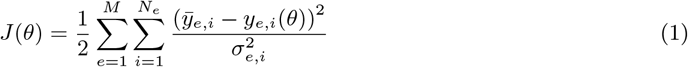

Here, *M* denotes the number of different experimental conditions, *N_e_* the number of measured data points for condition *e*, *y̅_e,i_* are the measured data points, *y_e,i_* are the observables from the model simulation, and *σ_e,i_* denotes the standard deviation for data point *y̅_e,i_*. An experimental condition is the setting of a specific biological perturbation experiment, such as a stimulation with a drug and its dosage, but it can also comprise differences in the experiment which are due to working with different cell-lines. In the ODE model, these experimental settings are given by a vector of input parameters, which define the initial value problem and hence determine the time-evolution of the studied system. If the system has *M* experimental conditions, this means that the underlying ODE model must be evaluated *M* times for different input parameter vectors, i.e., *M* different initial value problems have to be solved to evaluate the (full) objective function. A more detailed explanation of this aspect is given in the Methods section, an explanation in a more general context is given in the Supplementary Information.

Classical (full-batch) optimization methods evaluate the full objective function, i.e., simulate all experimental conditions, in each iteration of parameter optimization (Fig 1A). In contrast, mini-batch optimization methods evaluate only the contribution to the objective function coming from a randomly chosen subset, a mini-batch, of experimental conditions in each step (19; 49). The cycle until the whole dataset has been simulated, i.e., the computational equivalent of one iteration in full-batch optimization, is called an epoch. It is important to note that each experimental condition is simulated only once per epoch, i.e., the experimental conditions are drawn in a random, but non-redundant fashion. In this way, mini-batch optimization allows to perform more – but less informed – optimization steps than classical full-batch approaches in the same computation time.

**Figure 1:**
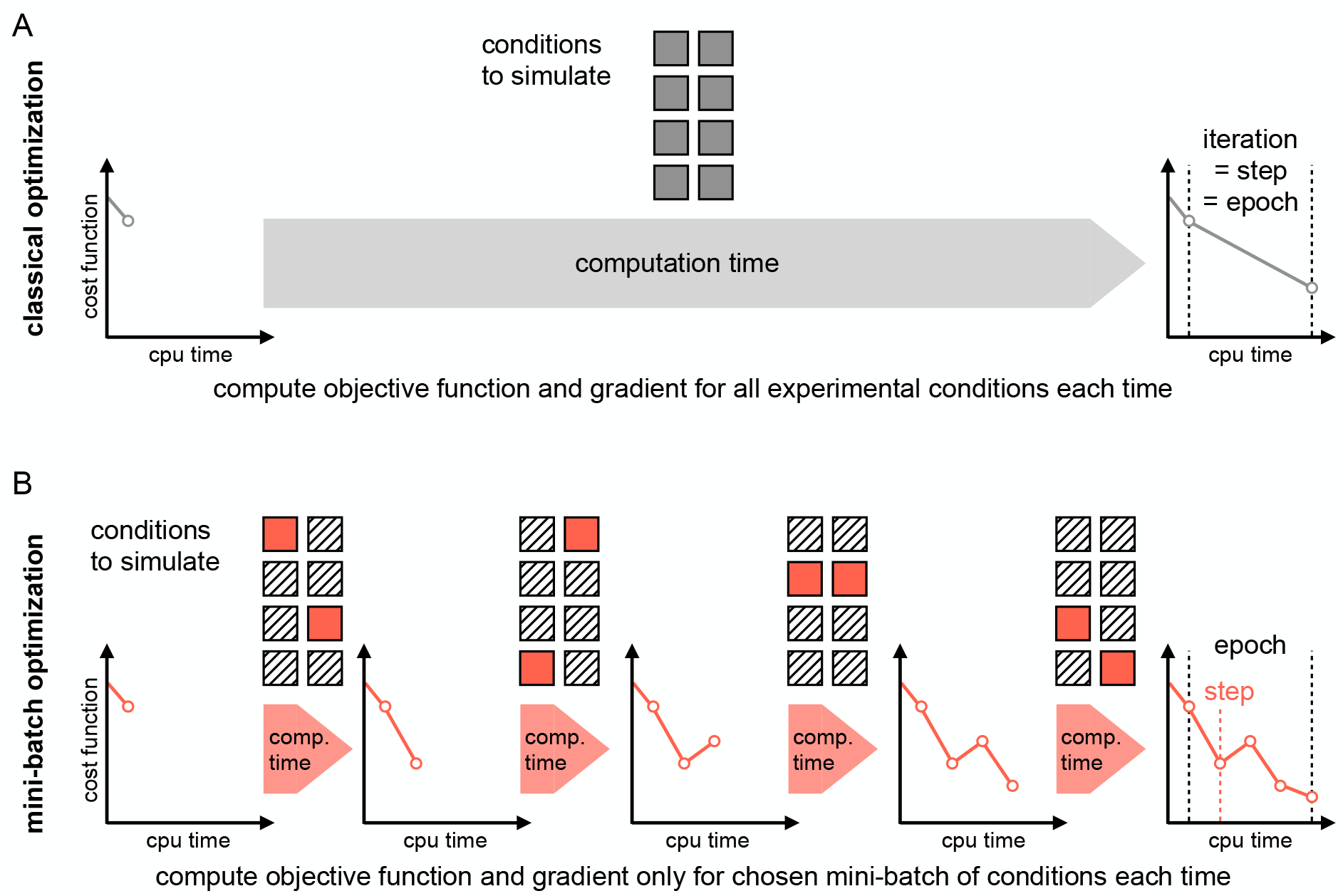
Visualization of full-batch and mini-batch optimization. **A** Classical full-batch optimization methods evaluate the contribution of all data points – and thus all experimental conditions – to the objective function in each step. The computation time scales linearly with the number of independently evaluable experimental conditions. **B** In mini-batch optimization, the independent experimental conditions are randomly divided into disjoint subsets, the mini-batches. Per optimization step, only the contribution of the chosen mini-batch is evaluated. Hence, possibly many optimization steps can be performed during one epoch, which is the time needed to evaluate the objective function on the whole dataset.

Various algorithms exist for full-batch and mini-batch optimization and each algorithm is influenced by different hyperparameters and optimizer settings. For full-batch optimization methods such as BFGS (18) and interior-point algorithms (64), many hyperparameters are associated with stopping conditions and at least good rules-of-thumb exist for their choice. For mini-batch optimization, there are various critical and less studied hyperparameters, e.g., the learning rate, which controls – but is not identical to – the size of the optimization step in parameter space. In order to apply mini-batch optimization methods to ODE models and benchmark the influence of these hyperparameters, we implemented some of the most common algorithms in the parallelizable optimization framework parPE (54): Vanilla stochastic gradient descent (SGD) (49), stochastic gradient descent with momentum (47; 58), RMSProp (61), and Adam (28) (see also Supplementary Information, Algorithms 1, 2, 3, and 4). This allowed a direct comparison with the implemented full-batch optimizers when using multi-start local optimization. More importantly, our implementation in parPE combines state-of-the-art numerical integration methods available in the SUNDIALS solver package (24) and adjoint sensitivity analysis for scalable gradient evaluation (13), since simple schemes (such as Euler’s method) can not be expected to yield reliable results for this problem class.

### Mini-batch size and learning rate schedules have a strong influence on optimizer performance

To evaluate available mini-batch optimization algorithms for ODE models, we considered three benchmark problems (Table 1, adapted from (20)). To facilitate the analysis of the scaling behaviour with respect to the number of experimental conditions, we generated artificial data (Fig. 2A). Details on the three benchmark examples and on the artificial datasets are given in the Methods section.

**Table 1.** Overview of ODE models for benchmarking mini-batch optimization

| Model name | State variables | Parameters | Conditions | Data points | Reference |
| --- | --- | --- | --- | --- | --- |
| Fujita | 9 | 19 | 600 | 6,000 | (17) |
| Bachmann | 25 | 40 | 1,200 | 12,000 | (3) |
| Lucarelli | 33 | 72 | 1,500 | 60,000 | (38) |

**Figure 2:**
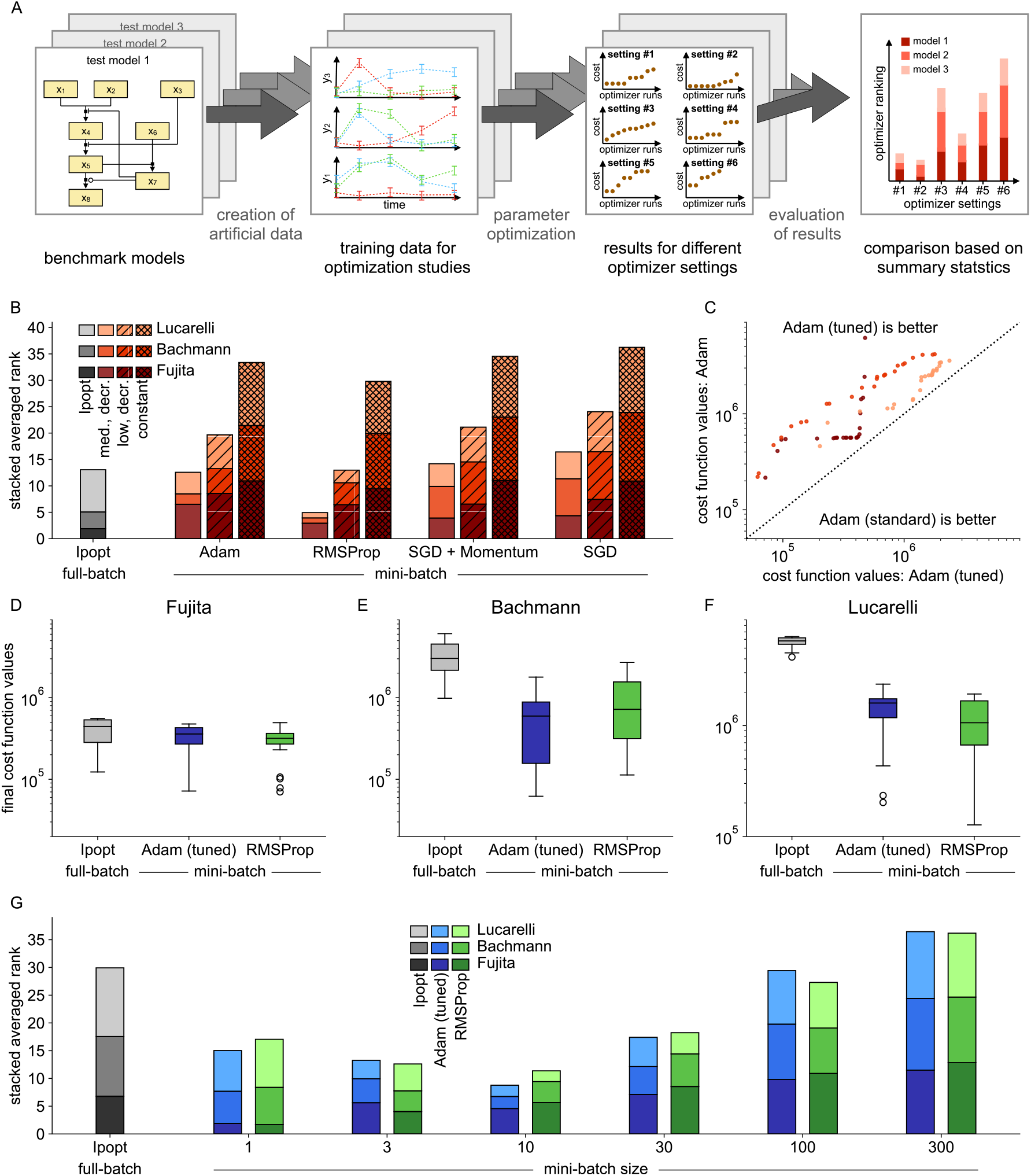
Bechmarking full-batch against mini-batch optimization methods on small- to medium-scale models. **A** Schematic overview of optimizer comparison: Benchmark models were chosen, noisy artificial data created, 100 initial points randomly sampled and different local optimizers started, each start was ranked between optimizers and an averaged score was computed. **B** Comparison of performance for different local optimizers with three different learning rate schedules (lower rank implies better performance, ranks of models are stacked). **C** Top 25 starts of the local optimizer Adam with tuning parameters taken from the literature (standard) vs. a simplified version (tuned). **D-F** Boxplots of final cost function values for the best 25 starts of the two best mini-batch optimizers, compared against the Ipopt (full-batch optimizer), for each model. **G** Comparison of performance for all starts of the best two mini-batch optimizers given the best learning rate, for different mini-batch sizes, compared against Ipopt (ranks of models are stacked).

We used a mini-batch size of 30 experimental conditions and 50 epochs of training – corresponding to roughly 50 iterations of a classical full-batch optimizer – which are typical hyperparameter choices in deep learning (19). We benchmarked the four implemented optimization algorithms: Vanilla stochastic gradient descent (SGD), stochastic gradient descent with momentum, RMSProp, and Adam (details are given in the Mathods section). To assess the impact of the learning rate, we considered three schedules: a medium and a low learning rate, both logarithmically decreasing, and a fixed learning rate. Details on these choices are given in the Methods section. The well-established full-batch optimizer Ipopt (64) was used as benchmark and was granted 50 iterations, so all tested methods had a similar computational budget. For each model, 100 randomly chosen initial parameter vectors were created, from which all optimizers were started. To assess the overall performance of each optimizer setting, we sorted the starts by their final objective function value and each of the 100 starts was ranked across the optimizer settings. Computing the mean of the 100 rankings for each setting led to an averaged rank, which we used as a proxy for overall optimization quality (Fig. 2A).

We found across all algorithms that the highest, i.e., the medium, but decreasing learning rate was preferred, the low but decreasing learning rate was second and the constant, medium learning rate resulted in the worst performance (Fig. 2B). A higher learning rate in the beginning of the optimization process seemed to be crucial for the mini-batch optimizers to progress quickly towards favorable regions of the parameter space (Supplementary Fig. 1, 2, and 3). Given the medium learning rate, different algorithms were able to compete with or even outperform the full-batch optimizer Ipopt, but the adaptive algorithm RMSProp performed particularly well. In most cases, the preferred learning rates led to step-sizes during optimization which were comparable or slightly lower than those which were chosen by classical (full-batch) optimization methods (Supplementary Fig. 4).

Given these findings, we compared the optimization algorithm Adam – which is maybe the most popular algorithm for training deep neural nets – with two different tuning variants: the tuning proposed in the original publication (called standard, see (28)) and a simplified scheme (called tuned), which employs the same rate for both internally used decaying averages (see Methods for more details). The analysis of the best 25 starts for all models with medium, decreasing learning rate showed that the tuned version outperformed the original one for all cases on our benchmark examples (Fig. 2C, Supplementary Fig. 1, 2, and 3). When comparing the performance of the tuned version of Adam and RMSProp with medium learning rate, we see that they show a very similar performance for the best 25 starts for all three tested models and perform as good as Ipopt or even better (Fig. 2D-F).

We then assessed the impact of the mini-batch size on the optimization result. Again, we used an average ranking, 100 starts, and investigated 6 mini-batch sizes for each model. We restricted our analysis to the two previously best performing optimization algorithms, tuned Adam and RMSProp, with the medium but decreasing learning rate. We found that in general, small mini-batch sizes were preferred, but the optimal size seemed to be model dependent (Fig. 2G, Supplementary Fig. 5, 6, and 7). While a mini-batch size of only one experimental condition worked best for the smallest example (Fujita), a mini-batch size of 10 experimental conditions performed best for the other two examples, yielding about 0.1% to 1% of the whole dataset as mini-batch. Interestingly, the mini-batch size seemed to impact both optimization algorithms to the same degree.

### Combining mini-batch optimization with backtracking line-search improves the robustness of the optimization process

A common challenge when performing parameter estimation for ODE models are regions in parameter space for which the numerical integration of the ODE is difficult or even fails. This may happen due to bad numerical conditioning of the problem or simply the divergence of the solutions (24). For this reason, full-batch optimizers use line-search or trust-region approaches (45), which can deal with these non-evaluable points by adapting the step-size (Fig. 3A). We found these problems also present in our benchmark examples (Fig. 3B, left), leading to failure of local optimization processes as available mini-batch optimization methods cannot handle failures of the objective function evaluation (probably because it is not encountered in deep learning). Hence, we implemented a rescue functionality, which attempts to recover a local optimization by undoing the previous step and performing backtracking line-search. In some cases, these failures happened at the initial points of optimization, and could hence not be recovered. In all of the remaining cases, the rescue functionality was able to successfully recover the respective local optimization (Fig. 3B). More details are given in the Methods section and Supplementary Information, Algorithm 5.

**Figure 3:**
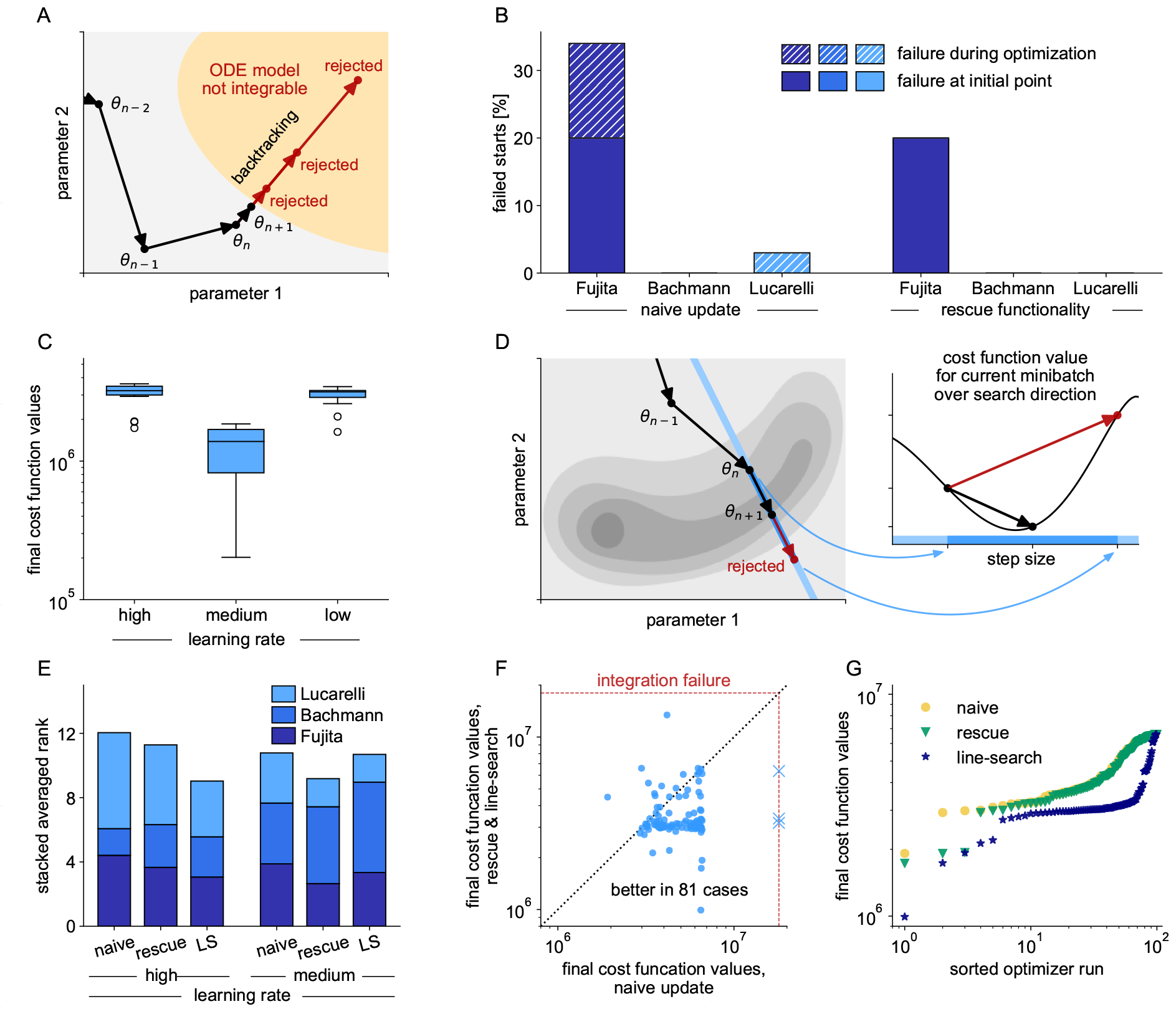
Influence of line-search methods on optimizers performance and reliability. **A** Schematic of the rescue functionality, which tries to recover from failed model evaluations, based on backtracking line-search. **B** Percentage of failed local optimizations per model (with optimizer Adam) due to non-integrability of the underlying ODE. Failure at the initial point of optimization can not be recovered, but failure during the optimization process is prevented when applying the rescue functionality. **C** Boxplots for the best 25 starts of mini-batch optimizers Adam for three different learning rates for the largest example (Lucarelli), showing that too high learning rates obstruct the optimization process. **D** Line-search implementation for mini-batch optimizers is implemented based on backtracking, while keeping the mini-batch fixed during line-search. **E** Comparison of performance for optimizer Adam, given different learning rates, for naive implementation, with rescue functionality, and rescue functionality and line-search (ranks of models are stacked). **F** All starts of the local optimizer Adam for the largest of the three examples (Lucarelli), naive implementation compared against rescue functionality and line-search, employing a high learning rate. **G** Waterfall plot for the largest of the three examples (Lucarelli), for naive implementation of Adam, with rescue functionality and with rescue functionality and line-search.

In the previous study, best optimization performance was achieved with medium learning rates. In the following, we increased the learning rate to higher values, but found it obstructing the optimization process (Fig. 3C). As overall, higher learning rates were beneficial and as it is a priori not clear for a given model what a good learning rate would be, we implemented an additional backtracking line-search.

It re-evaluates the objective function without gradient on the same mini-batch for different step-sizes, before accepting a proposed step (Fig. 3D). Details on the implementation can be found in the Methods section (Fig. 7) and Supplementary Information, Algorithm 6.

We evaluated these two algorithmic improvements for Adam and the medium and high learning rate on the three benchmark models (Fig. 3E, Supplementary Fig. 8). Interestingly, we found the strongest improvement for the largest model, although it suffers only little from integration failure (Fig. 3B). The line-search substantially improved the optimization process at high learning rates, which can be seen in a direct comparison (Fig. 3F) and in the waterfall plot (Fig. 3G). Considering all three models, we saw that the rescue functionality was generally helpful, whereas the line-search could also reduce the computational efficiency in case a good learning rate was chosen (Fig. 3E). This is not surprising, as the line-search needs additional computation time and some optimization runs were stopped prematurely due to imposed wall-time limits. However, these negative effects at lower learning rates were mild when compared against the positive effects at high learning rates and as the selection of a good learning rate is currently a trial-and-error process, the adaptation is highly beneficial.

### Mini-batch optimization enables training of predictive models of the drug response of cancer cell-lines

Following the successful testing and improvement, we evaluated how mini-batch optimization combines with sophisticated numerical integration algorithms applied for large-scale problems. Therefore, we considered the largest publicly available ODE model of cancer signaling (14). The model comprises various pathways and their cross-talk and captures 1,228 biochemical species and 2,686 reactions and was originally developed and provided by Alacris Theranostics. The generic chemical reaction network can be adapted to cancer cell-lines and treatment conditions using input parameter vectors. These vectors encode mutation and expression status (based on genome and transcriptome sequencing) and drug concentrations (Fig. 4A).

**Figure 4:**
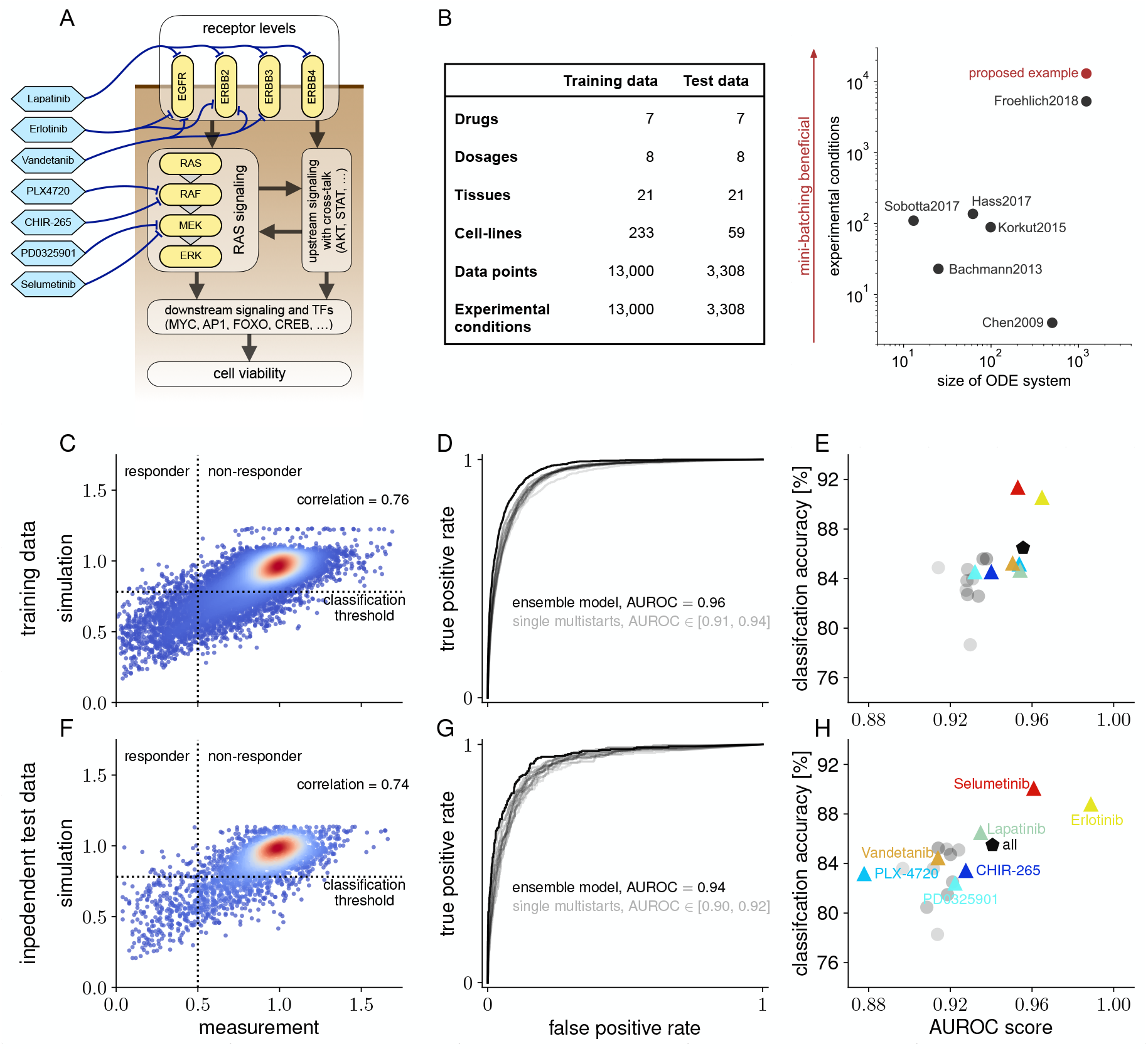
Description of application example, datasets and model performance, when trained with mini-batch optimization. **A** Simplifying illustration of the multi-pathway model of cancer signaling. **B** Overview of the datasets used for training and model validation, taken from the Cancer Cell Line Encyclopedia, and comparison of model sizes and experimental conditions used for model training of recently published ODE models. **C** Correlation of measured data and model simulation for all points of the training data, color-coding indicates density in scatter plot. **D** ROC-curves for classification into responsive and non-responsive combinations of cell-lines and treatments on training data for the best ten optimization runs (gray) and an ensemble simulation (black). **E** Area under ROC and classification accuracy on training data for the ten best optimization results (gray), for the ensemble model (black), and for the ensemble model on data for each drug individually (colored). **F** Correlation of measured data and model simulation for independent test data, color-coding indicates density in scatter plot. **G** ROC-curves for classification on test data. Classification thresholds from the training data were used. **H** Area under ROC and classification accuracy on test data for the ten best optimization results (gray), for the ensemble model (black), and for the ensemble model on data for each drug individually (colored).

We extracted all available drug response data from the Cancer Cell Line Encyclopedia (4), which we could match to the model, yielding in total 16,308 data points of viability read-outs. We split the data 80:20 into a training set and an independent test set. The training data is taken from 21 tissues with 7 different mechanistic targeted drugs at 8 different concentrations, adding up to 13,000 data points and experimental conditions (Fig. 4B). The test data comprises the same number of drugs and concentrations and is taken from 59 cell-lines from 21 (partly different) tissues, yielding 3,308 data points and experimental conditions. To the best of our knowledge, this is the first time that an ODE model has been trained on such a large set of training data from so many different experimental conditions.

We performed 100 local optimizations, in which we trained the model for 20 epochs and a mini-batch size of 100, using the optimization algorithm Adam (tuned) with rescue functionality as well as line-search. As in the Adam algorithm, the step-size during optimization scales with the square root of the problem dimension, we decided to take the medium learning rate schedule with a lower initial learning rate, yielding a step-size comparable to those for medium learning rates on the small- to medium-scale examples (details on these hyperparameter choices are given in the Methods section). We considered the best 10 optimization results for creation of an ensemble model, similar to (23). Based on this approach, we found a Pearson correlation of 0.76 of the simulation of the trained ensemble model with the training data (Fig 4C). We then considered a cell-line to be responsive to a particular treatment, if the viability of the corresponding cell-line was reduced by more than a factor of two. When computing the receiver-operator-characteristic (22) based on the trained ensemble model, we achieved an AUC value of 0.96 (Fig 4D). Interestingly, the AUC values when relying on single optimization runs instead of an ensemble were lower (between 0.91 and 0.94). Based on the characteristics, we computed classification thresholds for the simulation, when a cell-line together with a treatment condition is to be classified as responsive and repeated the computations for each considered drug individually. On the training data, the ensemble model achieved a classification accuracy of 86% (Fig 4E). Simulated responses to the drugs Selumentinib and Erlotinib matched particularly well, yielding AUCs of 0.95 and 0.96, and classification accuracies of 91% each, respectively.

When considering the independent set of test data and comparing it to the simulation of the ensemble model, we still found a Pearson correlation of 0.74, and hence only a slight decrease in model performance (Fig 4F). This suggest that there is only little to no overfitting. Computing the receiver-operator-characteristics yielded an AUC value of 0.94 for the ensemble model and AUC values between 0.90 and 0.92 for the ten best local optimizations (Fig 4G). This indicates that the trained model generalizes well to unseen cell-lines. We computed the classification accuracy, relying on the thresholds inferred from the training data and found that 85% of all treatment conditions were classified correctly into responsive or non-responsive (Fig 4H). Again, the AUCs and the classification accuracies varied across the considered drug, with Selumentinib and Erlotinib performing best also on the test data, with AUCs of 0.96 and 0.99 and classification accuracies of 90% and 89%, respectively.

### Backtracking line-search improves optimizer performance on the large-scale ODE model of cancer signaling

We then used the large-scale application example to confirm our findings from the smaller models. We compared 20 epochs of mini-batch optimization with Adam and either the former medium or the new lower learning rate schedules, with and without line-search, always using the rescue functionality, which enabled substantial improvements for this model (Supplementary Fig. 9). As benchmark, we performed 150 iterations with Ipopt – which also employs a line-search algorithm – and restricted to 20 local optimization due to the high computation time. All optimizers were initialized with the same parameter vectors. We also took snapshots of optimization process with Ipopt at computation times which were as close as possible to those used by the mini-batch optimizations.

We found that in terms of final objective function values and correlation with the training data, mini-batch optimization at lower learning rates yielded slightly better results than the Ipopt runs with 150 iterations (Fig. 5A). In terms of total computation time, the mini-batch approach was faster by a factor of 4.1. Optimization with medium learning rates yielded inferior results in terms of objective function values and correlation with the data, but further reduced the overall computation time. Additional line-search markedly improved the optimization quality for this model at the medium learning rate, while increasing the computation time by less than 13%. For the runs with lower learning rate, the line-search had almost no effect. When assessing the fit to independent test data, we found again that mini-batch optimization with the lower learning rates showed similar or better results than Ipopt (Fig. 5B). As before for high learning rates, the optimization at medium learning rates was improved by line-search, but turned out to be inferior overall.

**Figure 5:**
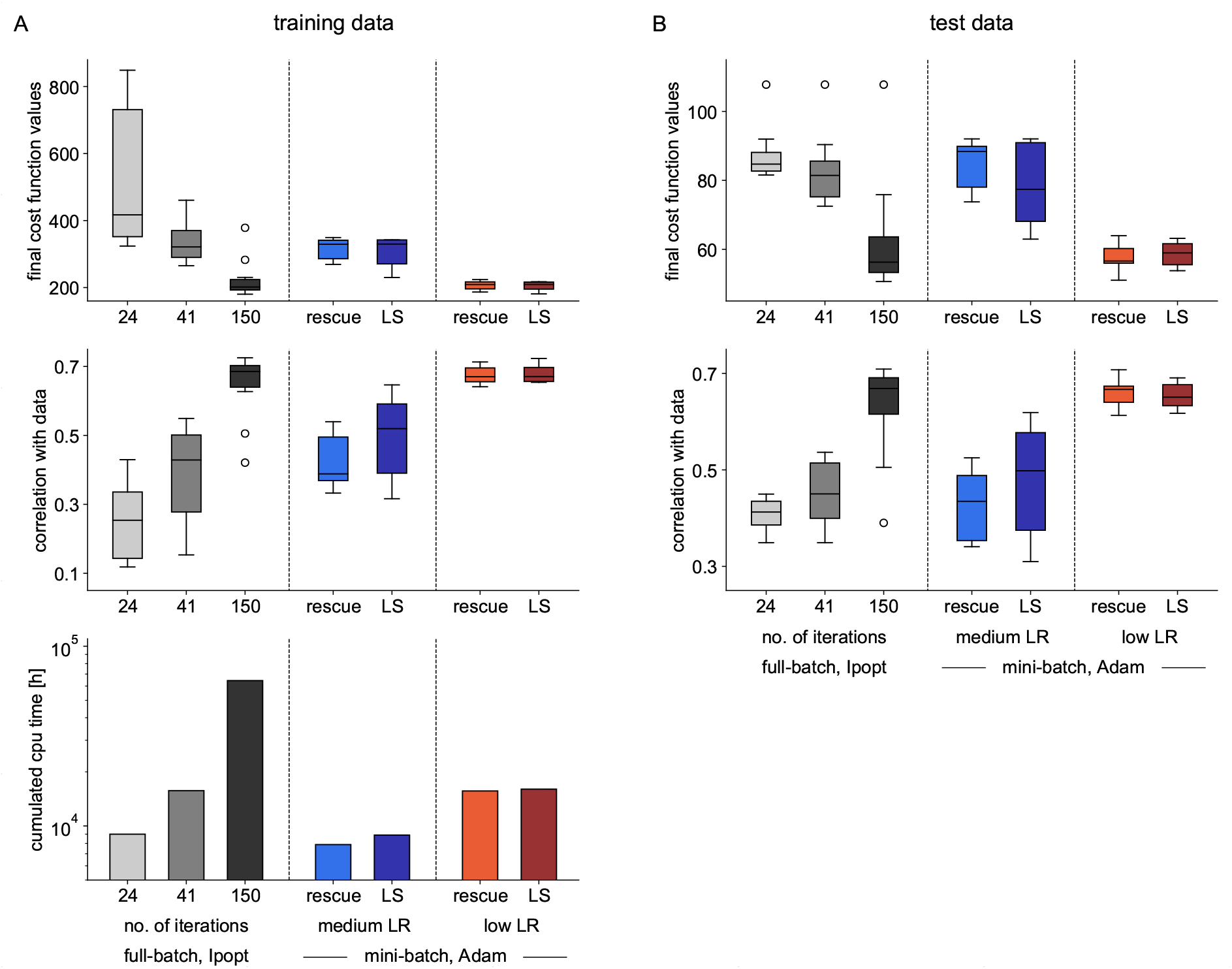
Comparison of optimization results for the large-scale cancer model, for different learning rates (LR), with rescue functionality only (rescue) and with additional line-search (LS). **A** Boxplots of the 10 best optimization runs out of 20, started at the same random parameters, for Ipopt (at three different stages of the optimization process to compare performance over computation time) and for mini-batch optimization with different optimization settings on training data: final objective functions values (upper panel), correlation of model simulation with measurement data (middle panel), and total computation time for all 20 optimization runs (lower panel). **B** Boxplots of the 10 best optimization runs, for Ipopt (at three different stages of the optimization process to compare performance over computation time) and for mini-batch optimization with different optimization settings on independent test data: final objective functions values (upper panel) and correlation of model simulation with measurement data (middle panel).

We investigated two further threshold-dependent characteristics to assess optimization performance: the computation time until convergence was reached for the first time and the number of converged starts per computation time. Both are common metrics for optimization performance (62). As threshold for convergence, we defined a value-to-reach based on the ten best optimization results from Ipopt and fixed it to the mean plus one standard deviation over final objective function values. We now granted 100 starts to the mini-batch optimizers, to allow them a similar budget of total computation time as for Ipopt. Considering the computation time until the first start converged, mini-batch optimization at medium learning rate with line-search was fastest, outperforming Ipopt by a factor of up to 27 (Supplementary Fig. 10). When comparing the number of converged starts per computation time, mini-batch optimization was up to a 6.9-fold faster than full-batch optimization. This time, optimization with lower learning rates performed better. Hence, lower learning rates yield more reasonable step-sizes for large-scale models. Overall, these observations confirm the finding that the choice of the learning rate is a crucial hyperparameter when working with mini-batch optimizers for ODE models and that, if too high learning rates are chosen, line-search can markedly improve results (Supplementary Fig. 10).

### Mini-batch optimization outperforms full-batch optimization by more than an order of magnitude

To evaluate the robustness of the mini-batch optimizer with respect to mini-batch size, we ran optimizations with mini-batch sizes 10, 100, 1000, and 13000 (full-batch), granting 10, 20, 50, and 150 epochs of optimization time and 100, 100, 50, and 25 local optimizations, respectively. This time, we used Adam without the line-search feature, as the previous study indicated it to have little to no impact for the chosen (low) learning rate.

For the large-scale model of cancer signaling, we found a clear benefit of smaller mini-batch sizes. Objective function and correlation values were substantially improved (Fig. 6A). As the total computation time was reduced by smaller mini-batch sizes, we faced the counter-intuitive effect of a seeming anti-correlation of optimization performance and total computation time. Optimization with the smallest mini-batch size outperformed Ipopt in terms of final objective function values while reducing the total computation time by more than a factor of 10. The findings on the optimization quality persisted when looking at the performance on independent test data (Fig. 6B). Again, the smallest mini-batch size yielded the best results, showing even a better generalization to independent test data than all previously tested approaches, possibly due the regularizing effect of small batch-sizes and hence possibly less overfitting of the training data.

**Figure 6:**
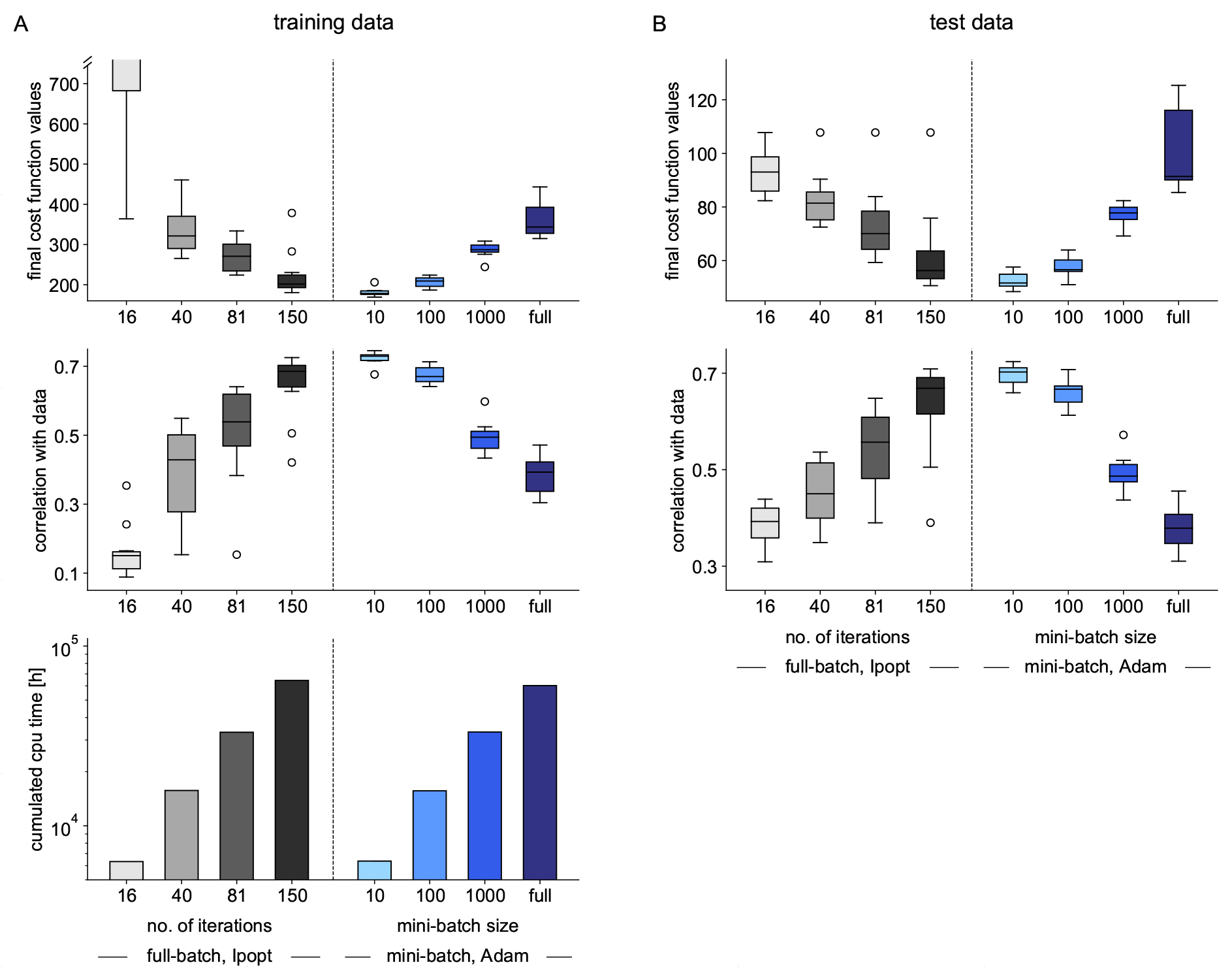
Comparison of optimization results for the large-scale cancer model, for different mini-batch sizes at low learning rate. **A** Boxplots of the 10 best optimization runs out of 20, started at the same random parameters, for Ipopt (at four different stages of the optimization process to compare performance over computation time) and for mini-batch optimization with different optimization settings on training data: final objective functions values (upper panel), correlation of model simulation with measurement data (middle panel), and total computation time for all 20 optimization runs (lower panel). **B** Evaluation of objective function values and correlation with measurement data for results from A on independent test data.

For the two smallest mini-batch sizes, we performed multi-start optimizations with 100 starts. Especially the waterfall plot for the smallest mini-batch size showed that not only computation time was reduced, but also the optimization quality was markedly improved (Supplementary Fig. 11). When comparing computation time to first convergence, mini-batch optimization was up to 52 times faster than Ipopt. In terms of converged starts per computation time, we found an 18-fold improvement when using mini-batch optimization. As additional test, we assessed the influence of the optimization algorithm on the optimization result (Supplementary Fig. 12). This indicated again that the chosen algorithm was less important than the choice of the learning rate or the mini-batch size.

## Discussion

We presented a framework for using mini-batch optimization in combination with advanced numerical integration methods for the parameter estimation of ODE models in systems biology. We introduced algorithmic improvements (tailored to ODE models), benchmarked different methods and their hyperparameters on published models (with artificial data), and identified the most important factors for successful optimization. Then, we applied mini-batch optimization to a particularly large personalizable model of cancer signaling and trained it on measured cell-line drug response data from a public database. The trained model provided accurate predictions whether a certain treatment would reduce cell viability by more than 50% for a chosen cell-line in more than 85% of the cases, even on cell-lines which had not been used for model training. The stochasticity introduced by the mini-batching in combination with the large dataset seemed to have addressed the overfitting observed in previous studies (14) and ensemble modeling improved the prediction accuracy further. Overall, our implementation reduced computation time by more than one order of magnitude while providing better fitting results than established methods. The improved scaling characteristics should render problems with even larger datasets feasible.

We identified the choices of learning rate and mini-batch size to be the most influential hyperparameters for parameter optimization and made the three following observations: Firstly, learning rates for mini-batch optimization which yield step-sizes slightly smaller than those used by established optimization techniques – see Supplementary Fig. 4 for examples – are a good choice. Secondly, surprisingly small mini-batch sizes were preferred in all of our application examples. Thirdly, the choice of the optimization algorithms seems to be less important, as at least Adam and RMSProp performed equally well on all examples.

Overall, learning rates and mini-batch sizes would be promising candidates for auto-tuning schemes. There are various known methods for auto-tuning of step-sizes during optimization (5; 6; 45). Combining those with mini-batch optimization may lead to substantial improvements and we also proposed and tested an implementation of a line-search method for mini-batching, which may serve as a starting point. For the mini-batch size, auto-tuning may be less straight forward, but also here, first approaches exist, which are based on assessing the variance of the objective function gradient across a chosen mini-batch and possibly enlarging the mini-batch size (34). Other algorithmic improvements – more specific to ODE models – would be combining mini-batch optimization with hierarchical optimization for observation-specific parameters, such as scaling factors or parameters for measurement noise (37; 54). This approach allows substantial improvements in parameter optimization for ODE models and it is to be expected that also mini-batch optimization would benefit from it. A complementary approach would be to implement variance reduction techniques (11; 53). Some of these methods enjoy good theoretical properties but are demanding in terms of memory consumption, which might make them prohibitive for applications in deep learning, but possibly well-suited for training of ODE models. Hierarchical optimization as well as variance reduction should be combined with methods for early-stopping, to avoid over-fitting and to further reduce computation time (39).

Since mini-batch optimization is computationally more efficient than full-batch optimization when working with large datasets, it is also a promising approach to drastically improve the exploration of parameter space. Especially when using methods such as multi-start local or hybrid local-global optimization (46; 48; 62), much more local optimizations can be performed. In our large-scale application example, we confirmed that ensemble modeling leads to better predictions than point estimates (9; 23). Furthermore, there have been recent advances when using ensembles created from the optimization history of an ODE model (63). Mini-batch optimization is particularly well-suited for these approaches, as it creates more comprehensive optimization histories.

In summary, we showed that combining mini-batch optimization with advanced numerical integration methods for parameter estimation of ODE models can help to overcome some major limitations. We hope it will become an actively developed and applied group of methods in systems biology. We think and hope that our work can serve as foundation for other research groups to further push the boundaries of what is computationally feasible and lead to new, fruitful applications.

## Methods

### Modeling of chemical reaction networks with ordinary differential equations (ODEs)

We considered ODE models with state vector 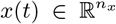, describing the concentrations of *n_x_* ∈ ℕ biochemical species, e.g., (phospho-)proteins or mRNA levels in a time interval *t* ∈ [0, *T*]. The time evolution of *x* was given by a vector field *f*, depending on unknown parameters 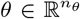, e.g., reaction rate constants, and a vector of known input parameters 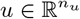:

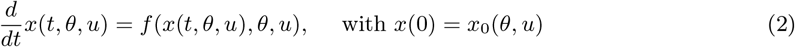

In our case, input parameters were drug treatments and differences between cell-lines based on mRNA expression levels and genetic profiles. As the ODE had no closed-form solution, we used numerical integration methods to solve/integrate Equation (2). As ODEs in systems biology applications are usually assumed to be stiff (30; 41), we employed an implicit multi-step backward differential formula scheme of variable order. This allowed adaptive time-stepping and automated error control, helping to ensure the desired accuracy of the computed results (30; 41; 48).

To match the model to the data, we used observable functions, which describe (phospho-)protein concentrations for the small- to medium-scale models. For the large-scale model, there is only one observable function (cell viability), described by a combination of downstream signaling activities (14):

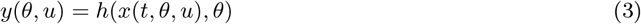

The data *D* = {*y̅_e,i_*}*e*=1,…,*M*, *i*=1,…,*N*_*e*_ was simulated as a time-course for the small- to medium-scale models. For the large-scale model, it was taken at steady-state, yielding only one datapoint per experimental condition *e*. Distinct experimental conditions differed through their vectors of input parameters *u_e_*, *e* = 1, …, *M*. Those input parameters captured all the differences between the different experimental setups, i.e., drug treatments and mRNA expression levels of the cell-lines. Hence, to simulate the whole dataset *D* once, *M* different initial value problems had to be solved.

To account for the fact that experimental data are noise-corrupted, we chose an additive Gaussian noise model with standard deviation *σ_e,i_* for experimental condition *e* and measurement index *i*. For the large-scale application example, we used the same *σ_e, i_* for all experiments, as no prior knowledge on the standard deviation was available.

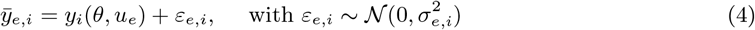

A more detailed explanation of ODE modeling in general is given in the Supplementary Information.

### Parameter optimization

This statistical observation model allowed us to compute the likelihood of an observed value *y*(*x*(*t*, *θ*, *u*), *θ*) given a parameter vector *θ*, assuming independence of the measurement noise terms (48). Due to its better numerical properties, we took its negative logarithm, which yielded:

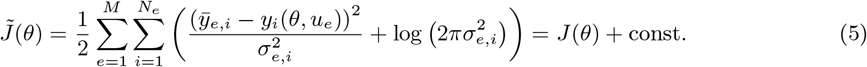

Assuming fixed measurement noise, the logarithmic term was just a constant offset. By neglecting it and identifying *y_i_*(*θ*, *u_e_*) with *y_e,i_*(*θ*), we arrived at the objective or cost function *J*(*θ*), which was given in Equation (1).

For global optimization of *θ*, we used multi-start local optimization, i.e., we randomly sampled many parameter vectors, from which we initialized local optimizations. This approach has repeatedly shown to be among the most competitive methods (48; 62), if high-performing local optimization methods with accurate gradient information of the objective function are used (52). In order to compute accurate gradients, we employed adjoint sensitivity analysis, which is currently the most performing method for gradient computation of high-dimensional ODE systems (13; 14). To benchmark our local optimization methods, we used the interior-point optimizer Ipopt (64), which combines a limited-memory BFGS scheme with a line-search approach (44). In previous studies (54), since such interior-point optimizers have shown to be among the most competitive methods for local optimization of large-scale ODE systems (54; 62).

More information on the formulation of the (log-)likelihood function and global parameter optimization of ODE models in a more general context is given in the Supplementary information.

### Mini-batch optimization algorithms

Mini-batch optimization is a particular type of local optimization, which exploits the sum structure of the objective function (49). In our case, we could rewrite the objective function in the following form:

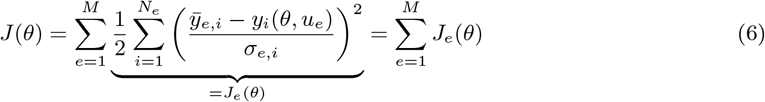

In the beginning of each epoch, the data set was randomly shuffled and then divided into mini-batches, random subsets of same size *S* ⊆ {1,…, *n*_*e*_}. Hence, no datapoint was used redundantly within one epoch. In each optimization step *r*, only the (sub-)gradient, sometimes also called gradient estimate (19), based on the mini-batch *S_r_* was used. The exact way how a parameter update was executed, i.e., how *θ*^(*r*+1)^ was computed from *θ*^(*r*)^ and the (sub-)gradient, 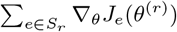 was dependent on the chosen algorithm. We investigated the following common mini-batch optimization algorithms in our study (see Supplementary Information for more details and (19; 50) for a comprehensive summary of mini-batch optimization algorithms):

- Vanilla stochastic gradient descent (SGD) (49), which is the simplest possible algorithm, using only the negative gradient of the objective function as update direction (Supplementary Information, Algorithm 1).
- Stochastic gradient descent with momentum (47; 58), a common variant, which uses a decaying average of negative gradients as direction instead of the negative objective function gradient alone (Supplementary Information, Algorithm 2).
- RMSProp (61), a so-called adaptive algorithm, which rescales/preconditions the current gradient by a decaying average over root-mean-squares of the previous objective function gradients (Supplementary Information, Algorithm 3).
- Adam (28), another adaptive algorithm, which attempts to combine the benefits of RMSProp with the momentum approach by using two decaying averages (Supplementary Information, Algorithm 4).

For Adam, we tested two different settings: As the two decaying averages in the algorithm are controlled by two tuning parameters *ρ*_1_ and *ρ*_2_, we set them first – according to the original publication – to 0.9 and 0.999, respectively, and then, based on some non-exhaustive testing, both to 0.9. We denoted the first setting as Adam (standard), the second as Adam (tuned).

### Learning rates and optimizer step-sizes

All the considered mini-batch algorithms rescale the computed parameter update with a factor called learning rate *η*, which can either be fixed over the optimization process, preschuled, or adapted according to the optimization process. In our study, we tested – based on the literature (19; 50) and our experience with local optimization – in total four learning rate schemes, which refer to the following numerical values for the small- to medium-scale models:

- High learning rate, logarithmically decreasing from 10^0^ to 10^−3^ (only used for studying the behavior when combined with line-search).
- Medium learning rate, logarithmically decreasing from 10^−1^ to 10^−4^.
- Low learning rate, logarithmically decreasing from 10^−2^ to 10^−5^.
- Constant learning rate, fixed to the value 10^−3^.

Assuming a given algorithm in optimization step *r* produced a parameter update (direction) *δ_r_*, then the next proposed parameter vector would be

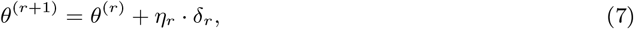

with *η_r_* being the learning rate at step *r*. Obviously, the learning rate influences the step-size of the optimization algorithm in parameter space. However, many of the algorithms we investigated yielded parameter updates with ‖*δ_r_*‖ ≠ 1. For, e.g., Adam, we obtained step sizes scaling with 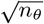, with *n_θ_* being the dimension of the unknown parameter vector (see Supplementary Information for more details and the corresponding calculation). When transferring the results of our study from the small- and medium-scale models to the large-scale model, we tried to conserve the actual step-sizes of the optimizers rather than the learning rates themselves, assuming the step-sizes to be the more fundamental quantities. On the large-scale model, we used two learning rate schedules with the following names and values:

- Medium learning rate, logarithmically decreasing from 10^−1^ to 10^−4^.
- Low learning rate, logarithmically decreasing from 10^−2^ to 10^−4^.

### Line-search and rescue functionality for mini-batch optimization

The rescue functionality was implemented as an iterative backtracking line-search algorithm, performing at most ten iterations. It is triggered, if the objective function and its gradient can not be evaluated. In this case, it keeps the current mini-batch, but undoes the previous parameter update, reducing the step length in each line-search iteration. More details on the method and its pseudo-code are given in the Supplementary Information (Algorithm 5 for the pseudo-code).

The additional line-search functionality was implemented according to the interpolation method, described in Chapter 3 of (45), and limited to at most three iterations. In each optimization step, the objective function value is checked on the same mini-batch after the parameter update. The parameter update is accepted, if the objective function decreases, otherwise, the step-size is reduced and the update repeated (Fig. 7). More details on the implementation and its pseudo-code are given in the Supplementary Information (Algorithm 6 for the pseudo-code).

**Figure 7:**
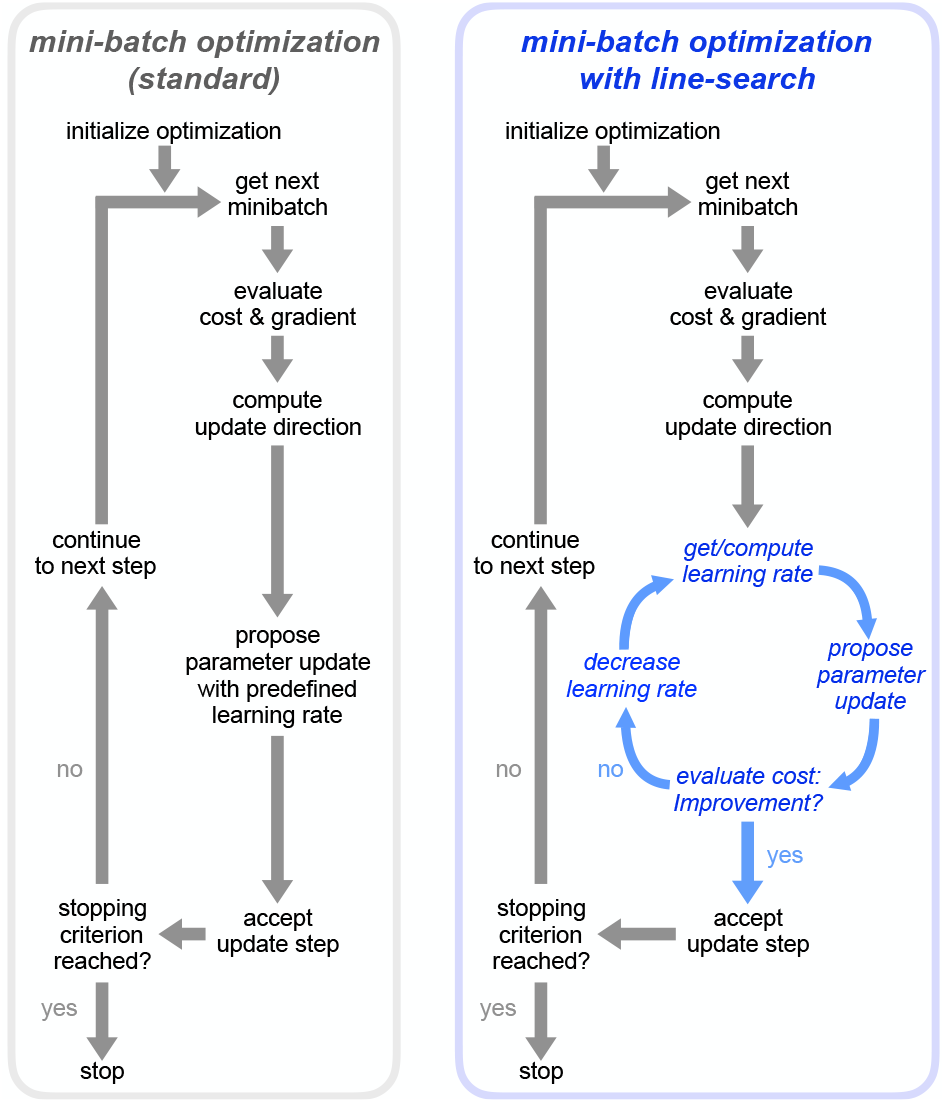
Comparison of standard mini-batch optimization and mini-batch optimization with line-search. Left panel: Standard mini-batch optimization uses a prescheduled learning rate, which determines the step-size during optimization regardless of whether an optimization step leads to an improvement or not. Right panel: If line-search is enabled, the objective function is re-evaluated on the same mini-batch and checked for improvement. If no improvement is achieved, the learning rate is reduced until either improvement is achieved or until the maximum number of line-search steps is reached.

### Computation of final objective function and correlation values

To ensure an unbiased comparison of objective values, we computed final objective function and correlation values after optimization for all methods (full-batch and mini-batch) on the whole dataset. As we performed this comparison also on a set of independent test data and since the model was using scaling parameters to match the model output to the measurement data (48), we computed those scaling parameters analytically (54; 65).

### Computation of receiver-operator-characteristics and classification accuracy

We used 13,000 datapoints from 233 cell-lines for model training and 3,308 datapoints from 59 cell-lines as test set. However, only for 198 cell-lines from the training set and 49 cell-lines from the test set, all treatment conditions were available. In order to compute unbiased receiver-operator-characteristics (ROCs), we used only those cell-lines, yielding 11,088 datapoints for the training set and 2,744 datapoints for the test set. Experimental conditions were grouped into two groups: Those, in which cell viability was reduced by more than 50% when compared to the untreated condition, were defined as responsive, the rest as non-responsive. We then computed classification thresholds to be those model output values, which corresponded to the points on the ROC being tangential to an affine function with slope 1 ≙ 45°.

### Implementation of parameter estimation using the toolboxes AMICI and parPE

Parameter estimation was performed using the parPE C++ library (54), which provides the means for parallelized objective function evaluation and optimization of ODE models generated by the AMICI ODE-solver toolbox (16) using optimizers such as Ipopt (64). In our studies, we used Ipopt version 3.12.9, running with linear solver ma27 and L-BFGS approximation of the Hessian matrix and extended parPE with the mini-batch algorithms described above.

For numerical integration of the ODEs we used AMICI, which provides a high-level interface to the CVODES solver (55) from the SUNDIALS package (24) and generates model specific C++ code for model evaluation and likelihood computation to ensure computational efficiency. In our applications, we used AMICI default settings with adjoint sensitivitity analysis, employing a backward differential formula (BDF) scheme of variable order, with adaptive time stepping and error tolerances of 10^−8^ for the relative and 10^−16^ for the absolute integration error per step, allowing at most 10^4^ integration steps.

Optimizations were run on the SuperMUC phase 2 supercomputer (Leibniz Supercomputing Centre, Garching, Germany). Compute nodes were equipped with two Haswell Xeon Processor E5-2697 v3 CPUs (28 cores per node) and 64GB of RAM. For the large-scale model, mini-batch optimization multi-starts comprising 100 local optimizations were run on 65 nodes (1820 cores) with a wall-time limit of 48 hours. The 20 local optimizations of Ipopt were separated to single runs with 12 nodes (336 cores), and 35 hours of wall-time were granted. For the small to medium-scale models, each of the multi-starts comprising 100 local optimizations was run on 1, 2, and 3 nodes for the Fujita, Bachmann, and the Lucarelli model, respectively, always exploiting all 28 cores per node. Wall-times were fixed to 15, 32, and 40 hours, respectively.

### Adaptation of benchmark models and creation of artificial data

The small- to medium-scale examples for the benchmark study were chosen based on a collection of benchmark models (20). We chose models with different system sizes, which allowed the generation of large artificial datasets, which were sufficiently heterogeneous. This should ensure clear differences in objective function values and gradients, when different mini-batches were used. In order to allow the creation of heterogeneous datasets, the SBML files and input parameters were slightly altered. The precise model versions are made freely available in the SBML/PEtab format at Zenodo, under https://doi.org/10.5281/zenodo.3556429 (59).

Artificial data was created by simulating the models with the parameter vectors reported in (20). Additive Gaussian noise was added to the model simulations, using the noise levels which were reported in (20) for each observable.

## Data and Code availability

The employed models, datasets, and codes for parameter estimation and subsequent analysis are freely available at Zenodo, under https://doi.org/10.5281/zenodo.3556429 (59) in the code versions, which were used to generate the results. We refer users interested in using our implementations to the respective Github pages, where more recent versions of the toolboxes are made freely available: https://github.com/ICB-DCM/parPE/ (for the parallelized parameter estimation code, including the implementation of the mini-batch algorithms), https://github.com/ICB-DCM/AMICI/ (for the implementation of the ODE solver, likelihood and gradient computation using adjoint sansitivity analysis), and https://github.com/ICB-DCM/PEtab/ (for the model and data format of the small to medium-scale examples). The SBML-model of the large-scale application example is available at https://github.com/ICB-DCM/CS_Signalling_ERBB_RAS_AKT, the data is provided as hdf5 file.

## Acknowledgements

This work was supported by the European Union’s Horizon 2020 research and innovation program under grant agreement no. 686282 (B.L., C.W., D.W., J.H., P.S.), CanPathPro. Computer resources for this project have been provided by the Gauss Centre for Supercomputing / Leibniz Supercomputing Centre under grant pr62li.

## Author information

### Author contributions

D.W. designed and implemented the parallelized optimization framework, P.S. implemented the mini-batch optimization algorithms, designed and implemented their algorithmic improvements. C.W. and B.L. retrieved and processed the data. J.H. and P.S. conceived the computational studies for mini-batch optimization, P.S., L.S., J.H. and D.W. conceived the studies for full-batch optimization. P.S. and L.S. performed the studies. P.S., L.S., J.H. and D.W. analyzed the results. All authors wrote and approved the final manuscript.

