## Supplementary information for "Mini-batch optimization enables training of ODE models on large-scale datasets"

November 2019

#### Parameter estimation for ordinary differential equation models

##### Modeling of chemical reaction networks with ordinary differential equations (ODEs)

Ordinary differential equations (ODEs) are a common approach to model the time-evolution of bio-chemical species. The underlying assumptions when using ODE models are firstly, that spatial inhomogeneities can be neglected within one compartment and secondly, that all species are abundant enough that the process is governed by deterministic dynamics, in contrast to stochastic dynamics, which can be found in systems with low copy numbers.

In most applications, an ODE model describes the time evolution of a state vector $x(t) \in \mathbb{R}^{n_x}$ of concentrations of $n_x \in \mathbb{N}$ biochemical species, which are, for example, (phospho-)proteins or mRNA levels, at time $t \in [0, T]$. The time evolution of $x$ is given by a vector field $f$, which depends on unknown parameters $\theta \in \mathbb{R}^{n_\theta}$ and known input parameters $u \in \mathbb{R}^{n_u}$:

$$\frac{d}{dt}x(t, \theta, u) = f(x(t, \theta, u), \theta, u), \quad \text{with } x(0) = x_0(\theta, u) \tag{1}$$

Parameters are typically reaction rates constants or initial concentrations, which can not be measured. Input parameters are often external stimuli such as treatments with drugs or other ligands, or measurable initial concentrations, mRNA expression levels or genetic profiles. Depending on $f$, $\theta$, and $u$, a solution of the ODE given in Equation (1) may or may not exist for a time interval $[0, T]$. However, in most of the considered cases, it exists, but is not available in closed form. This makes it necessary to use numerical solvers to integrate/solve the ODE. Again depending on the equation, the employed ODE solver, and its settings, a

*To whom correspondence should be addressed.

numerical integration of Equation (1) may fail, which is a common problem when using ODE models of biological process, at least during parameter estimation (6; 7). Hence, involved implementations of, e.g., implicit Runge-Kutta, Adams-Moulton, or BDF schemes are used, which allow adaptive time-stepping and automated error control, to attempt to avoid integration failure and to ensure the desired accuracy of the computed results (6; 7; 10).

An ODE model also allows the description of the time-evolution of a vector of measurable quantities $y(t) \in \mathbb{R}^{n_y}$, which we will call observables. In realistic modeling applications, it is often impossible to measure state variables directly, as – in many cases – measurement techniques only allow the assessment on a relative scale (e.g., quantitative PCR), maybe with an additional offset (e.g., immunoblotting), on a (bi-)exponential scale (e.g., flow cytometry), or as sums of certain state-variables (e.g., if an antibody binds to a protein, independent of its phosphorylation state). For this reason, we introduce observable functions:

$$y(t, \theta, u) = h(x(t, \theta, u), \theta) \tag{2}$$

An observable can be matched to experimental data $D$, taken at timepoints $t_i, i = 1, \ldots, n_t$ and measured for $n_e$ experimental conditions. Distinct experimental conditions differ through the vectors of input parameters $u_k \in \mathbb{R}^{n_u}$, $k = 1, \ldots, n_e$, which describe all the differences in the model, which are necessary to capture different experimental setups, such as drug treatments or other external stimuli, growth conditions for bacteria or differences in, e.g., mRNA expression levels for different cell-lines. In most cases, $u$ determines the initial values of the ODE, but it can also comprise a time-resolved inputs, such as splines. To simulate the whole dataset $D$ once, $n_e$ different initial value problems have to be solved. $D$ is then given as:

$$D = \{\bar{y}_{ij}^k\}_{\substack{i=1,\ldots,n_y^k \\ j=1,\ldots,n_t}}^{k=1,\ldots,n_e} \tag{3}$$

Experimental data are noise-corrupted, and thus we have to choose a noise model. In this study, we follow the most common approach and assume additive Gaussian noise, yielding:

$$\bar{y}_{ij}^k = y_i(t_j, \theta, u^k) + \varepsilon_{ij}^k, \quad \text{with } \varepsilon \sim \mathcal{N}(0, \left(\sigma_{ij}^k\right)^2) \tag{4}$$

Here, $\sigma_{ij}^k$ denotes the standard deviations for the respective timepoints. This statistical model allows us to compute the likelihood of a measurement value $y_i(t_k, \theta, u^k)$ given a parameter vector $\theta$, assuming independence of the measurement noise terms:

$$\mathcal{L}(D \mid \theta) = \prod_{k=1}^{n_e} \prod_{i=1}^{n_y^k} \prod_{j=1}^{n_t} \frac{1}{\sqrt{2\pi}\sigma_{ij}^k} \exp\left\{ -\frac{1}{2} \left( \frac{\bar{y}_{ij}^k - y_i(t_k, \theta, u^k)}{\sigma_{ij}^k} \right)^2 \right\} \tag{5}$$

##### Parameter estimation, local, and global optimization

The statistical model makes it possible to fit the unknown model parameters to the measurement data, by maximizing the likelihood $\mathcal{L}(D \mid \theta)$. However, it is more common to work with the negative logarithm of the likelihood, as firstly, most optimization algorithms are implemented as minimization algorithms and secondly, the logarithm of the likelihood tends to be more convex and is hence numerically better tractable (10). This yields the negative log-likelihood as objective or cost function:

$$J(\theta) = -\log\left(\mathcal{L}(D \mid \theta)\right) = \frac{1}{2} \sum_{k=1}^{n_e} \sum_{i=1}^{n_y^k} \sum_{j=1}^{n_t} \left( \left( \frac{\bar{y}_{ij}^k - y_i(t_k, \theta, u^k)}{\sigma_{ij}^k} \right)^2 + \log\left(2\pi \left(\sigma_{ij}^k\right)^2\right) \right) \tag{6}$$

For additive Gaussian noise with known measurement noise $\sigma_{ij}^k$ – as assumed in this study – the logarithmic term is just a constant, offsetting the objective function value from 0, but not affecting the location of the minimum. In this case, minimizing this objective function is equivalent to using a weighted least squares algorithm. In parameter estimation/optimization, the aim is to find the global minimizer of $J$, which is the maximum likelihood estimator $\theta^{MLE}$ of $\mathcal{L}(D \mid \theta)$:

$$\theta^{MLE} = \text{argmin}_{\theta \in \Omega} J(\theta) \tag{7}$$

Hence, minimizing $J$ is equivalent to optimizing the quality of the fit of the model to the data. Usually, the parameter vector $\theta$ is restricted to a region $\Omega \in \mathbb{R}^{n_\theta}$ in parameter space, which is assumed to be biologically plausible.

For global optimization of $\theta$, many methods exist (14). A simple idea, called multi-start local optimization, is to run algorithms which are guaranteed to find a local minimum from many random initial points (10; 18). These results can be sorted by the final objective function values and visualized in waterfall plots, which can – if optimization worked well – reveal the structure of the local minima of the objective function. The best local optimum is then assumed to be the global optimum. Other common methods to find global optima are genetic or swarm-based algorithms, simulated annealing, or scatter-search, which combines genetic algorithms with multi-start local optimization. For ODE models however, it is well established that multi-start local optimization and scatter-search outperform other approaches for most, though maybe not for all, cases (10; 14; 18).

Local optimization algorithms usually exploit derivative information, such as the gradient of the objective function. There are three common methods of gradient computation for ODE models: finite difference schemes, forward sensitivity analysis, and adjoint sensitivity analysis. For large-scale ODE models with many parameters, adjoint sensitivity analysis (1) has been shown to be the most efficient approach and subsequently for local optimization (1; 2; 18). Different local optimization algorithms exist, among which gradient descent, quasi-Newton methods such as (L-)BFGS, Gauss-Newton, trust-region approaches such as the dogleg method, or interior-point methods are the most popular. For parameter estimation of very large ODE models, implementations of interior-point methods combined with (L-)BFGS schemes and either trust-region or line-search methods for controlling the step size have shown their effectiveness (1; 2; 15; 18).

### Mini-batch optimization

Mini-batch optimization is a particular type of local optimization and only capable of finding local optima (12). The basic idea consists of exploiting the fact that the objective function and its gradient have the structure of a sum:

$$J(\theta) = \frac{1}{2} \sum_{k=1}^{n_e} \underbrace{\sum_{i=1}^{n_y^k} \sum_{j=1}^{n_t} \left( \left( \frac{\bar{y}_{ij}^k - y_i(t_k, \theta, u^k)}{\sigma_{ij}^k} \right)^2 + \log\left(2\pi \left(\sigma_{ij}^k\right)^2\right) \right)}_{=J^k(\theta)} = \sum_{k=1}^{n_e} J^k(\theta) \tag{8}$$

$$\nabla_\theta J(\theta) = \ldots = \sum_{k=1}^{n_e} \nabla_\theta J^k(\theta) \tag{9}$$

In each optimization step $r$, a random subset – the mini-batch – $S_r \subseteq \{1, \ldots, n_e\}$ is chosen, and the objective function and its gradient are computed only based on $S_r$. It is important to note that no experimental condition is chosen twice until the full dataset has been evaluated, a time-frame which is called one epoch. Hence, the mini-batches belonging to one epoch are disjoint. Theoretically, also the inner sum structure (over the indices $i$ and $j$) could be used for mini-batching. However, when working with ODE models, the computational demanding part is the solution of the initial value problem for an experimental condition $k$. Once this is done, evaluating the different observables $y_i$ at different time points $t_j$ happens at negligible cost. Hence, for most applications on ODE models, mini-batching will only make sense over the conditions $k$.

##### Mini-batch optimization algorithms in a nutshell

The way how the parameter update is executed, i.e., how $\theta^{(r+1)}$ is computed from $\theta^{(r)}$ and the (sub-)gradient $\sum_{k \in S_r} \nabla_\theta J_k(\theta^{(r)})$, depends on the chosen algorithm. Some of the most commonly used algorithms which we investigated in our study are:

- Vanilla stochastic gradient descent (SGD) (12), which is the simplest possible algorithm, using only the negative gradient of the objective function as update direction (Supplementary Information, Algorithm 1).
- Stochastic gradient descent with momentum (9; 16), a common variant, which uses a decaying average of negative gradients as direction instead of the negative objective function gradient alone (Supplementary Information, Algorithm 2).
- RMSProp (17), a so-called adaptive algorithm, which rescales the current gradient by a decaying average over root-mean-squares over the previous objective function gradients (Supplementary Information, Algorithm 3).
- Adam (5), another adaptive algorithm, which attempts to combine the benefits of RMSProp with the momentum approach by using two decaying averages (Supplementary Information, Algorithm 4).

A good summary of these and further mini-batch optimization algorithms can be found in (3).

As Adam is a very popular algorithm and has more tuning parameters than the other algorithms, we investigated two different settings of these tuning parameters: The two decaying averages are governed by two coefficients, usually denoted as $\rho_1$ and $\rho_2$, averaging the gradient and its norm, respectively. The original publication sets the former to a value of 0.9 and the latter to 0.999, which we denoted as standard version of Adam. We also investigated a simplified/tuned version, in which we set $\rho_1 = \rho_2 = 0.9$.

**Algorithm 1** Parameter update for vanilla stochastic gradient descent

*Initialization*:

**Set** initial learning rate $\eta_0$
**Set** final learning rate $\eta_N$
**Set** number of epochs $N$

*Procedure for parameter update during optimization*:

**Get** index of current epoch $k$
**Get** current parameter vector: $\theta$
**Get** gradient estimate for current mini-batch: $g$
Compute update direction (normalized negative gradient estimate): $\delta \leftarrow -g/\|g\|$
Compute current learning rate: $\eta \leftarrow \eta(k, \eta_0, \eta_N, N)$
Update parameter vector: $\theta \leftarrow \theta + \eta \cdot \delta$

##### Learning rates and step-sizes during optimization

Despite their different approaches, all the mentioned mini-batch algorithms have some features in common. Firstly, they are not guaranteed to – and, in fact, don't even try to – produce a series of parameter vectors, along which the objective function decreases monotonically. Hence, it is to be expected that the progress

**Algorithm 2** Parameter update for stochastic gradient descent with vanilla momentum

*Initialization*:

**Set** initial learning rate $\eta_0$
**Set** final learning rate $\eta_N$
**Set** number of epochs $N$
**Set** decay rate for momentum $\alpha = 0.8$
**Set** momentum vector $v = 0 \in \mathbb{R}^{n_\theta}$

*Procedure for parameter update during optimization*:

**Get** index of current epoch $k$
**Get** current parameter vector: $\theta$
**Get** gradient estimate for current mini-batch: $g$
**Get** momentum vector: $v$
Compute new momentum: $v \leftarrow \alpha v + (1 - \alpha) g$
Compute update direction: $\delta \leftarrow -v / \max(\|v\|, 1)$
Compute current learning rate: $\eta \leftarrow \eta(k, \eta_0, \eta_N, N)$
Update parameter vector: $\theta \leftarrow \theta + \eta \cdot \delta$

**Algorithm 3** Parameter update step for RMSProp

*Initialization*:

**Set** initial learning rate $\eta_0$
**Set** final learning rate $\eta_N$
**Set** number of epochs $N$
**Set** decay rate for gradient norm history $\rho = 0.9$
**Set** gradient norm history vector $h = 0 \in \mathbb{R}^{n_\theta}$
**Set** stabilization constant $\varepsilon = 10^{-7}$

*Procedure for parameter update during optimization*:

**Get** index of current epoch $k$
**Get** current parameter vector: $\theta$
**Get** gradient estimate for current mini-batch: $g$
**Get** gradient norm history vector: $h$
Compute new gradient norm history vector (using element-wise multiplication): $h \leftarrow \rho h + (1 - \rho) g \odot g$
Compute update direction (using element-wise division): $\delta \leftarrow -g \oslash (\sqrt{h} + \varepsilon)$
Compute current learning rate: $\eta \leftarrow \eta(k, \eta_0, \eta_N, N)$
Update parameter vector: $\theta \leftarrow \theta + \eta \cdot \delta$

**Algorithm 4** Parameter update step for Adam

*Initialization*:

**Set** initial learning rate $\eta_0$
**Set** final learning rate $\eta_N$
**Set** number of epochs $N$
**Set** decay rate for gradient history $\rho_1 = 0.9$
**Set** decay rate for gradient norm history $\rho_2 = 0.999$ (or $\rho_2 = 0.9$ for tuned version)
**Set** gradient history vector $h_1 = 0 \in \mathbb{R}^{n_\theta}$
**Set** gradient norm history vector $h_2 = 0 \in \mathbb{R}^{n_\theta}$
**Set** stabilization constant $\varepsilon = 10^{-7}$

*Procedure for parameter update during optimization*:

**Require:** Index of current optimization step $\ell$

**Get** index of current epoch $k$
**Get** current parameter vector: $\theta$
**Get** gradient estimate for current mini-batch: $g$
**Get** gradient history vector: $h_1$
**Get** gradient norm history vector: $h_2$
Compute new gradient history vector: $h_1 \leftarrow \rho_1 h_1 + (1 - \rho_1) g$
Compute new gradient norm history vector: $h_2 \leftarrow \rho_2 h_2 + (1 - \rho_2) g \odot g$
Compute update direction (using element-wise division, applying bias-correction):
$\delta \leftarrow -\frac{h_1}{1-\rho_1^\ell} \oslash \left( \sqrt{\frac{h_2}{1-\rho_2^\ell}} + \varepsilon \right)$
Compute current learning rate: $\eta \leftarrow \eta(k, \eta_0, \eta_N, N)$
Update parameter vector: $\theta \leftarrow \theta + \eta \cdot \delta$

which is made in one optimization step can be (at least partly) undone in another, subsequent step. Over the whole optimization process however, most of them are guaranteed to converge to a local minimum in a probabilistic sense. This is a striking difference to most local full-batch optimization algorithms, which at least try to or can even guarantee to produce monotonically decreasing trajectories of objective function values. Secondly, all of these mini-batch optimization algorithms rescale the proposed optimization step with a factor $\eta$, called the learning rate. If we assume a given algorithm in optimization step $r$ to produce a parameter update (direction) $\delta_r$, then the next proposed parameter vector will be

$$\theta^{(r+1)} = \theta^{(r)} + \eta_r \cdot \delta_r, \tag{10}$$

with $\eta_r$ being the learning rate at step $r$. Obviously, $\eta$ affects the step size of the optimizer. But as for most algorithms $\|\delta_r\| \neq 1$, it is not identical to the step size. In many publications, the learning rate is either fixed to a previously chosen value, such as $10^{-3}$ (13), which is a commonly used value for training deep neural nets, or prescheduled over the optimization process (3). In our studies on the small- to medium-scale models, we worked in total with four different learning rates:

- A high learning rate, which logarithmically decreased from $10^0$ to $10^{-3}$ (only used for studying the behavior when combined with line-search).
- A medium learning rate, which logarithmically decreased from $10^{-1}$ to $10^{-4}$.
- A low learning, which logarithmically decreased from $10^{-2}$ to $10^{-5}$.
- A constant learning rate, fixed to the value $10^{-3}$.

We assumed that the crucial quantity for the optimization process is the actual optimizer step-size and not the learning rate. From our experience with parameter estimation of ODE models, we aimed at initial optimizer step-sizes to be roughly in the interval $[1, 10]$, rather tending towards 1 for smaller and rather tending towards 10 for larger models. It is important to note that this is a mere rule-of-thumb based on personal experience. However, this implies that different learning rates had to be chosen for the different models, since for Adam, the initial step size is given by the following computation:

$$\|\Delta\theta\| = \|\eta\delta\| = \sqrt{\sum_{j=1}^{n_\theta} \frac{\frac{(1-\rho_1)g_j}{1-\rho_1^1}}{\sqrt{\frac{(1-\rho_2)g_j^2}{1-\rho_2^1}} + \varepsilon}} = \sqrt{\sum_{j=1}^{n_\theta} \frac{g_j}{g_j + \varepsilon}} \approx \sqrt{n_\theta} \tag{11}$$

Here, we used the notation from Algorithm 6. As we achieved best results for medium learning rates on the small- and medium-scale models, we aimed at similar optimizer step-sizes on the large-scale model. This implied that we had to reduce the learning rates by an order of magnitude, as the large-scale model has about two order of magnitude more parameters than the small- and medium-scale models. For this reason, we used the two following learning rates with the two following names for the large-scale model:

- Medium learning rate, logarithmically decreasing from $10^{-1}$ to $10^{-4}$.
- Low learning rate, logarithmically decreasing from $10^{-2}$ to $10^{-4}$.

### Enhancements of mini-batch optimization for ODE models

##### Motivation from established methods used in full-batch optimization of ODE models

Classical full-batch local optimization algorithms, which are used for parameter estimation of ODE models, employ trust-region or line-search methods for adaptive step-sizing (4; 11). These approaches provide an intuitive way of dealing with integration failure of the underlying ODE: If the ODE can not be solved, the objective function value and its gradient are set to infinity. This makes the proposed step unacceptable and leads to a shrinkage of the step-size, until the ODE can be solved again. In mini-batch optimization, similar approaches do not exist, which complicates a direct method transfer to parameter optimization of ODE models. Hence, we introduced two functionalities to overcome this issue. In both of them, we adapt the current learning rate, by multiplying it with a reduction factor $\beta$, which mimics the role of a line-search or a one-dimensional trust-region.

##### Rescue functionality to avoid ODE integration failure

The rescue functionality is activated, if ODE integration (with gradient computation) fails: It undoes the last optimization step (but keeps the current mini-batch), multiplies the reduction factor $\beta$ by $r$ (in our implementation, we set $r = 0.2$, but any number $r < 1$ would do), and proposes a new step, for which the objective function and its gradient are computed. This procedure is repeated at most ten times, until either a parameter vector is found for which ODE integration is possible, or the local optimization run is finally stopped. Whenever ODE integration is successful, the reduction factor is mildly increased again by multiplication with $c$ (in our case: $c = 1.3$, but any number $c > 1$ would do), to at most 1. In our implementation, it takes about six steps until the old step-size is recovered after ODE-integration failure. A detailed pseudo-code of this functionality is given in Algorithm 5.

##### Additional line-search functionality to improve optimization for high learning rates

The additional line-search works independently of the rescue functionality and is based on the interpolation method, described in Chapter 3 of (8). After a parameter update is proposed, the objective function is evaluated without gradient – for the new parameter vector, but on the same mini-batch – and checked for improvement. If the new step yields an improvement, it is accepted, otherwise a backtracking line-search is performed according to the mentioned algorithm, by shrinking the reduction factor $\beta$. A comparison of this approach to standard mini-batch optimization is given in Fig. 7 of the main manuscript, a detailed pseudo-code is provided in Algorithm 6.

Rescue functionality and additional line-search can be combined, as they are not redundant. This can be seen in the following way: The additional line-search implementation may reduce step-sizes, if integration failure is encountered, but will finally accept one of the proposed steps after (at least in our study) three iterations. If the ODE can yet not be integrated, the rescue functionality will be activated in the next optimization step. It also occurs that the ODE can be integrated for a given parameter vector, but the numerical computation of its gradient is not possible. Also such a case will trigger the rescue functionality, despite an additional line-search being used.

**Algorithm 5** Rescue functionality for mini-batch optimization

Code parts specific to rescue functionality are shown in blue

*Initialization*:

**Set** initial parameter vector $\theta \leftarrow \theta_0$

**Set** and initialize optimization algorithm for parameter update

**Set** stopping criterion, e.g., maximum number of epochs $N$

**Set** variables to store necessary information about previous optimization step: parameter vector $\theta^*$, gradient estimate $g^*$, possibly optimization algorithm dependent quantities $Q^*$, initialize with $(\theta^*, g^*, Q^*) \leftarrow (\mathrm{NaN}, \mathrm{NaN}, \mathrm{NaN})$

**Set** reduction factor $\beta = 1$

**Set** reduction multiplier $r < 1$

**Set** increase multiplier $c > 1$

*Procedure for optimization with rescue functionality*:

**while** Stopping criterion not met **do**
    **Get** next mini-batch
    Compute gradient estimate: $g \leftarrow \mathrm{gradientFunction}(\theta)$
    **if** ODE integration was successful **then**
        Update old quantities: $(\theta^*, g^*, Q^*) \leftarrow (\theta, g, Q)$
        Update reduction factor: $\beta \leftarrow \min(c \cdot \beta, 1)$
        Call parameter updater: $\theta \leftarrow \mathrm{parameterUpdater}(\theta, g, Q, \beta)$
    **else**
        **if** $\theta^*$ is NaN **then**
            **return** ODE integration failure at initial point
        **end if**
        **while** Last call to gradientFunction failed **and** maximum number of rescue steps not reached **do**
            Undo last step: $(\theta, g, Q) \leftarrow (\theta^*, g^*, Q^*)$
            Update reduction factor $\beta \leftarrow r \cdot \beta$
            Call parameter updater: $\theta \leftarrow \mathrm{parameterUpdater}(\theta, g, Q, \beta)$
            Compute gradient estimate: $g \leftarrow \mathrm{gradientFunction}(\theta)$
        **end while**
        **if** Last call to gradientFunction failed **then**
            **return** ODE integration failure not recoverable
        **end if**
    **end if**
**end while**

*Changes in* parameterUpdater-*functions*:

Replace parameter update $(\theta \leftarrow \theta + \eta \cdot \delta)$ by $(\theta \leftarrow \theta + \eta \cdot \beta \cdot \delta)$

**Algorithm 6** Line-search functionality for mini-batch optimization
Code parts specific to line-search functionality are shown in blue

*Initialization*:

**Set** initial parameter vector $\theta \leftarrow \theta_0$
**Set** and initialize optimization algorithm for parameter update
**Set** stopping criterion, e.g., maximum number of epochs $N$
**Set** variable to store information from previous optimization step: parameter vector $\theta^*$

*Procedure for optimization with line-search functionality*:

**while** Stopping criterion not met **do**
  **Get** next mini-batch
  Compute objective and gradient estimate: $\left(j,g\right) \leftarrow$ gradientFunction($\theta$)
  **Set** line-search reduction factor $\gamma = 1$
  Call parameter updater: $\theta \leftarrow$ parameterUpdater($\theta, g, Q, \gamma$)
  Compute objective: $j^{(1)} \leftarrow$ objectiveFunction($\theta$)
  **if** Improvement: $j^{(1)} < j$ **then**
    Accept step: $\theta^* \leftarrow \theta$
  **else**
    Undo last step: $\theta \leftarrow \theta^*$
    Compute optimal line-search factor $\gamma \leftarrow$ based on quadratic interpolation for $g, j, j^{(1)}$
    Call parameter updater: $\theta \leftarrow$ parameterUpdater($\theta, g, Q, \gamma$)
    Compute objective: $j^{(2)} \leftarrow$ objectiveFunction($\theta$)
    **if** Improvement: $j^{(2)} < j$ **then**
      Accept step: $\theta^* \leftarrow \theta$
    **else**
      Undo last step: $\theta \leftarrow \theta^*$
      Compute optimal line-search factor $\gamma \leftarrow$ based on cubic interpolation for $g, j, j^{(1)}, j^{(2)}$
      Call parameter updater: $\theta \leftarrow$ parameterUpdater($\theta, g, Q, \gamma$)
      Compute objective: $j^{(3)} \leftarrow$ objectiveFunction($\theta$)
      **Set** $m = 3$
      **while** No improvement **and** maximum number of line-search steps not reached **do**
        Undo last step: $\theta \leftarrow \theta^*$
        Compute optimal line-search factor $\gamma \leftarrow$ based on cubic interpolation for $g, j, j^{(m-1)}, j^{(m)}$
        Call parameter updater: $\theta \leftarrow$ parameterUpdater($\theta, g, Q, \gamma$)
        Compute objective: $j^{(m+1)} \leftarrow$ objectiveFunction($\theta$)
        Increment $m \leftarrow m + 1$
      **end while**
      Accept step: $\theta^* \leftarrow \theta$
    **end if**
  **end if**
**end while**

*Changes in* parameterUpdater-*functions*:

Replace parameter update ($\theta \leftarrow \theta + \eta \cdot \delta$) by ($\theta \leftarrow \theta + \eta \cdot \gamma \cdot \delta$)

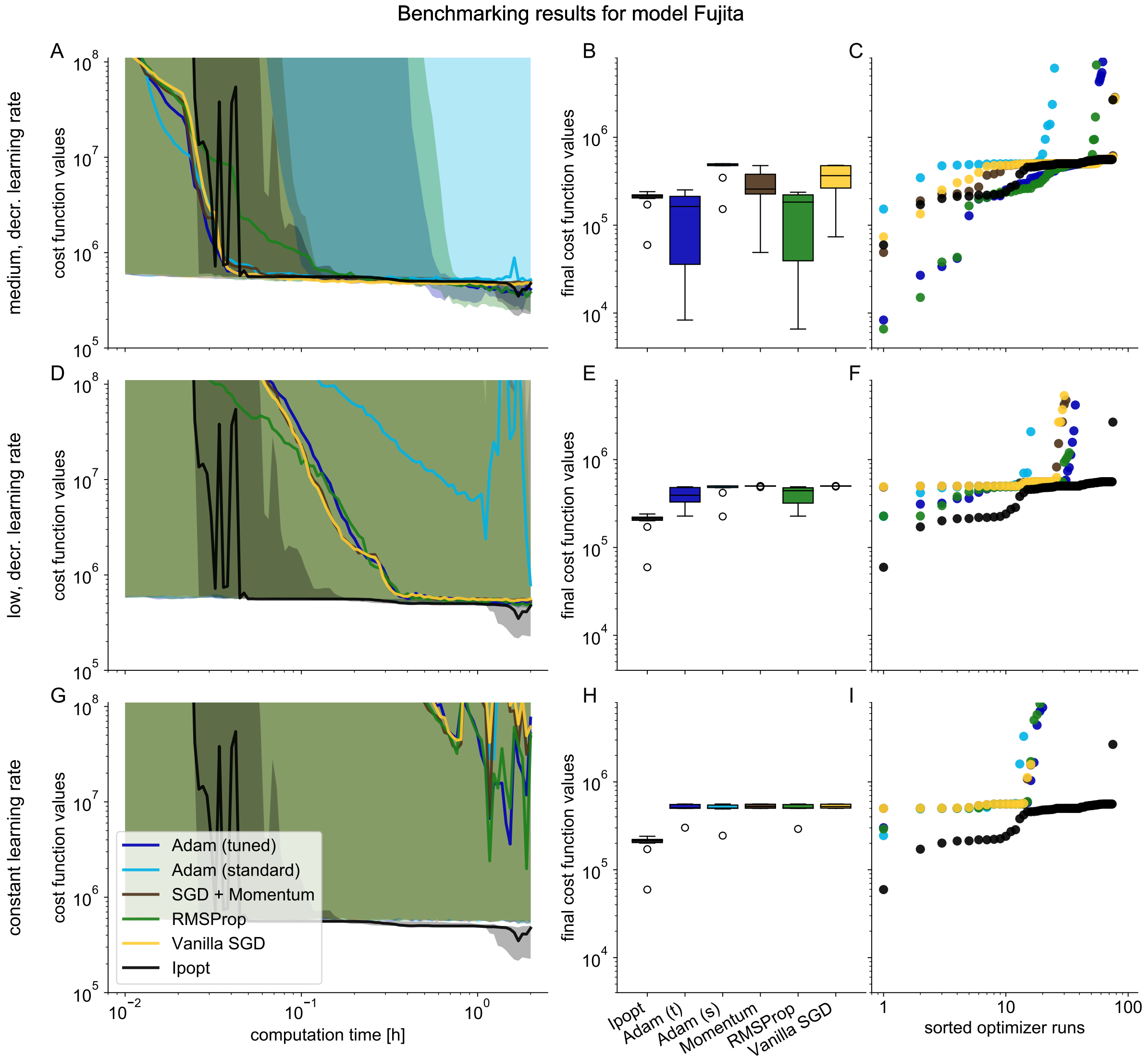


Figure 1: Medium learning rates show best performance for mini-batch optimization of ODE models (Supplementary to Fig. 2 of main manuscript). Different learning rate schemes for mini-batch optimization methods were benchmarked against full-batch optimizer Ipopt on Fujita example model. Objective function values along optimization traces were extrapolated from mini-batch to the full dataset, smoothed over multiple optimization steps to filter the stochasticity of the mini-batches, and interpolated for a mesh of computation time points. Shading depicts percentiles for 10% and 40% of optimizer traces, solid lines depict percentiles for 20%. Final objective function values for box and waterfall plots were evaluated at final point of optimization trajectory. **A** Traces of objective function value during optimization for medium, decreasing learning rate. **B** Box plots of the 10 best final objective function values for medium, decreasing learning rate. **C** Waterfall plot of all final objective function values for medium, decreasing learning rate. **D** Traces of objective function value during optimization for low, decreasing learning rate. **E** Box plots of the 10 best final objective function values for low, decreasing learning rate. **F** Waterfall plot of all final objective function values for low, decreasing learning rate. **G** Traces of objective function value during optimization for constant learning rate. **H** Box plots of the 10 best final objective function values for constant learning rate. **I** Waterfall plot of all final objective function values for constant learning rate.

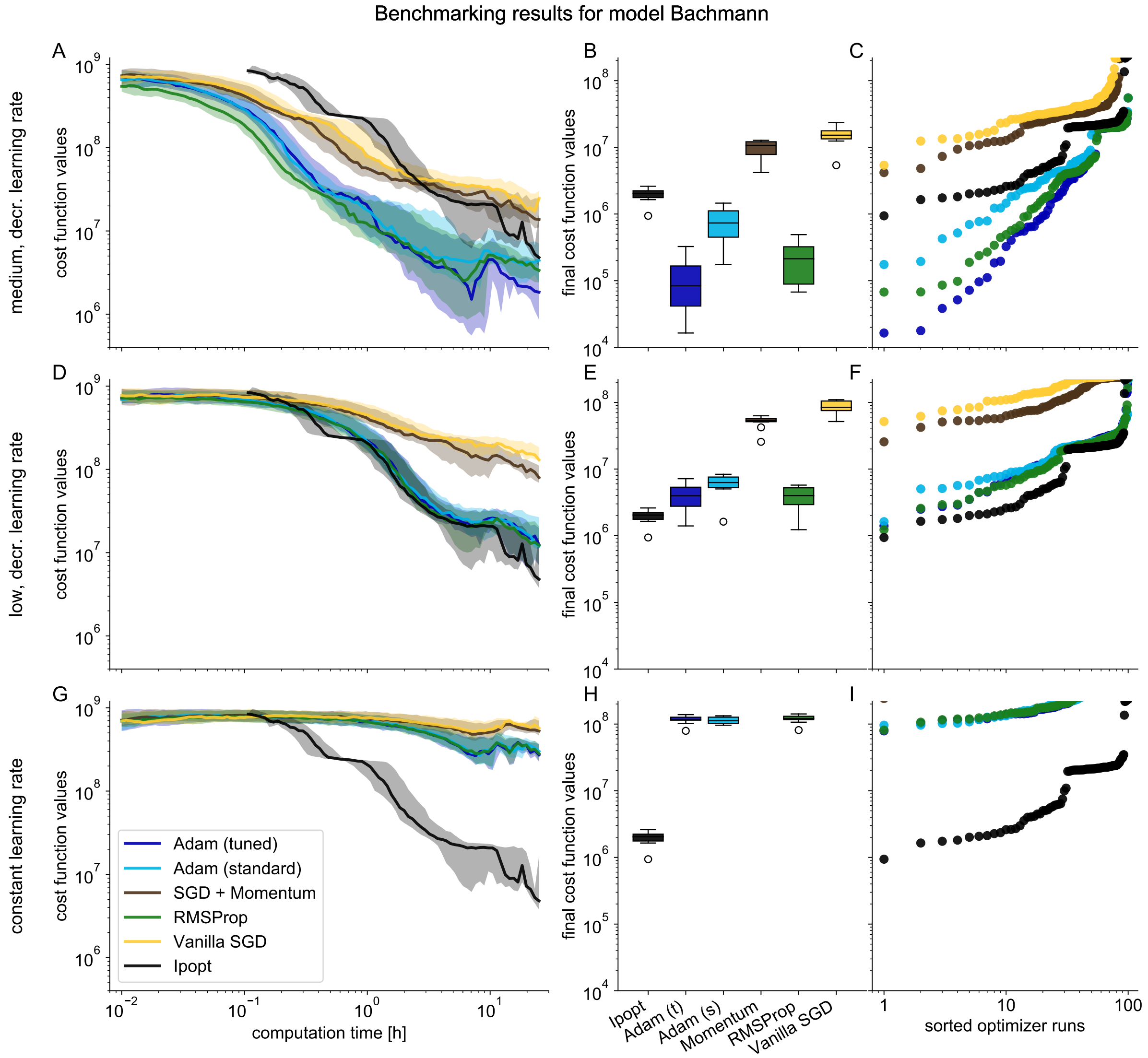


Figure 2: Medium learning rates show best performance for mini-batch optimization of ODE models (Supplementary to Fig. 2 of main manuscript). Different learning rate schemes for mini-batch optimization methods were benchmarked against full-batch optimizer Ipopt on Bachmann example model. Objective function values along optimization traces were extrapolated from mini-batch to the full dataset, smoothed over multiple optimization steps to filter the stochasticity of the mini-batches, and interpolated for a mesh of computation time points. Shading depicts percentiles for 10% and 40% of optimizer traces, solid lines depict percentiles for 20%. Final objective function values for box and waterfall plots were evaluated at final point of optimization trajectory. **A** Traces of objective function value during optimization for medium, decreasing learning rate. **B** Box plots of the 10 best final objective function values for medium, decreasing learning rate. **C** Waterfall plot of all final objective function values for medium, decreasing learning rate. **D** Traces of objective function value during optimization for low, decreasing learning rate. **E** Box plots of the 10 best final objective function values for low, decreasing learning rate. **F** Waterfall plot of all final objective function values for low, decreasing learning rate. **G** Traces of objective function value during optimization for constant learning rate. **H** Box plots of the 10 best final objective function values for constant learning rate. **I** Waterfall plot of all final objective function values for constant learning rate.

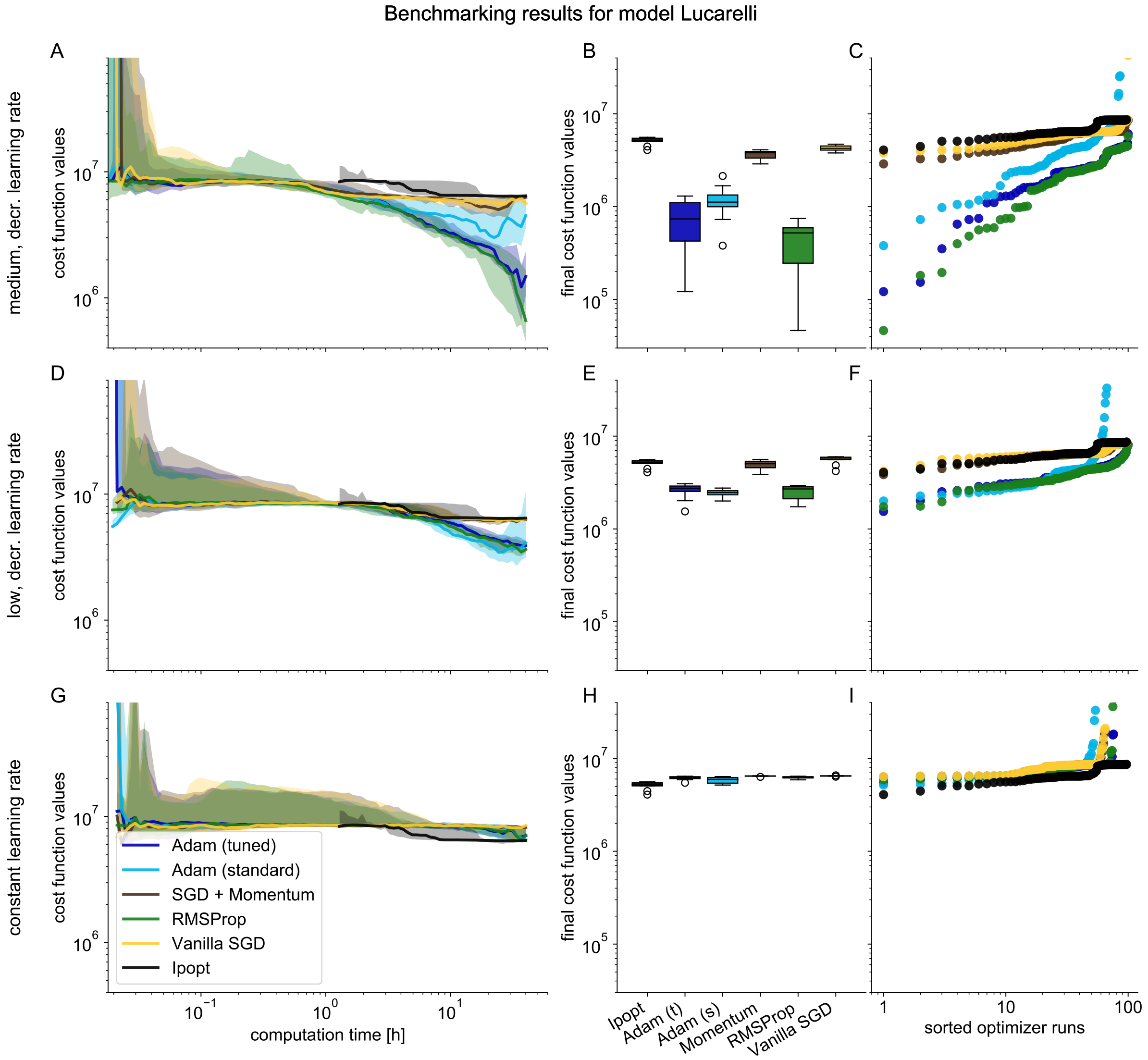


Figure 3: Medium learning rates show best performance for mini-batch optimization of ODE models (Supplementary to Fig. 2 of main manuscript). Different learning rate schemes for mini-batch optimization methods were benchmarked against full-batch optimizer Ipopt on Lucarelli example model. Objective function values along optimization traces were extrapolated from mini-batch to the full dataset, smoothed over multiple optimization steps to filter the stochasticity of the mini-batches, and interpolated for a mesh of computation time points. Shading depicts percentiles for 10% and 40% of optimizer traces, solid lines depict percentiles for 20%. Final objective function values for box and waterfall plots were evaluated at final point of optimization trajectory. **A** Traces of objective function value during optimization for medium, decreasing learning rate. **B** Box plots of the 10 best final objective function values for medium, decreasing learning rate. **C** Waterfall plot of all final objective function values for medium, decreasing learning rate. **D** Traces of objective function value during optimization for low, decreasing learning rate. **E** Box plots of the 10 best final objective function values for low, decreasing learning rate. **F** Waterfall plot of all final objective function values for low, decreasing learning rate. **G** Traces of objective function value during optimization for constant learning rate. **H** Box plots of the 10 best final objective function values for constant learning rate. **I** Waterfall plot of all final objective function values for constant learning rate.

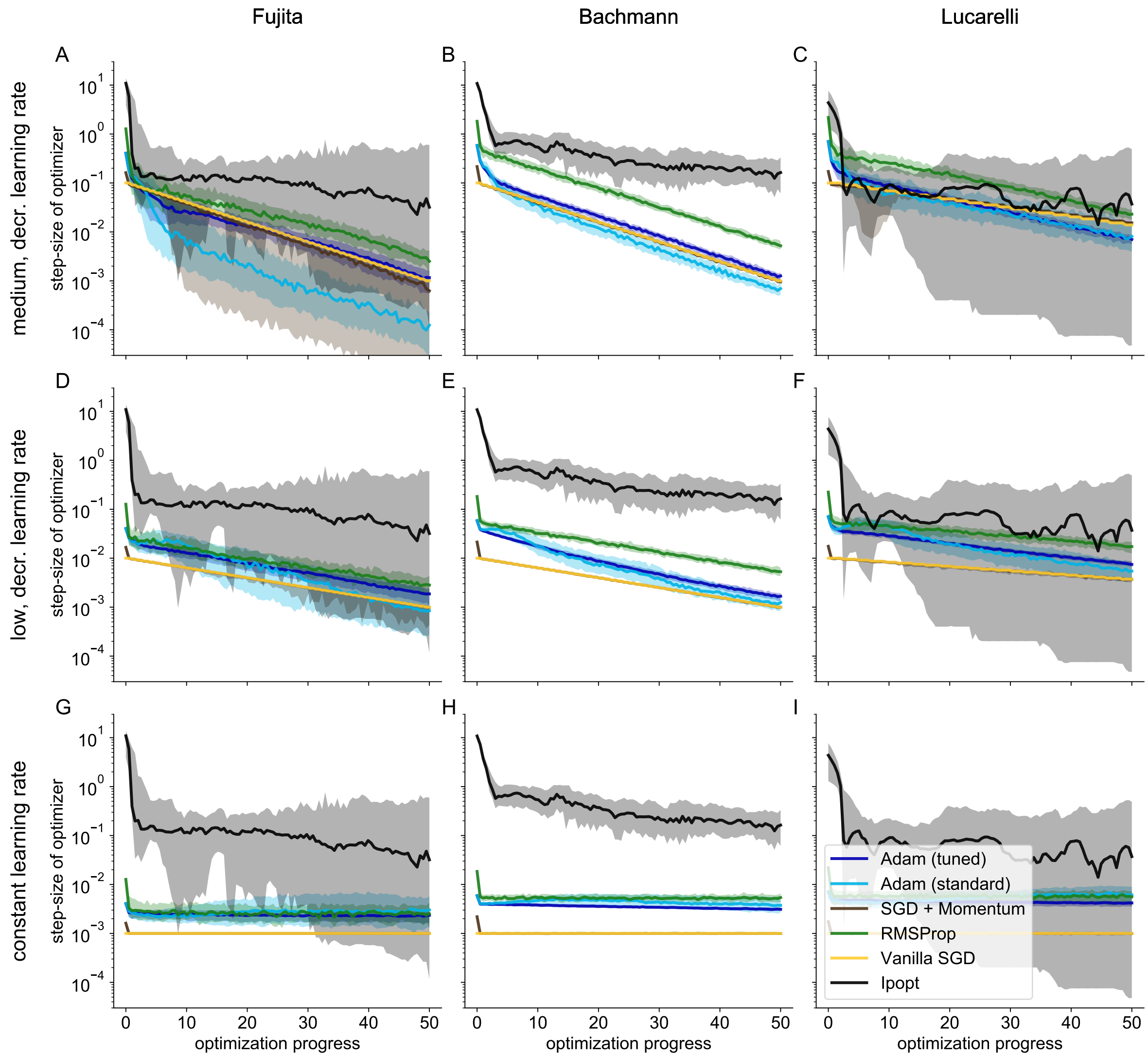


Figure 4: Learning rates leading to step-sizes slightly smaller than those of full-batch optimization methods perform best (Supplementary to Fig. 2 of main manuscript). Effective step-sizes of optimization steps in parameter space are plotted for full-batch optimizer Ipopt and different mini-batch optimization algorithms, against optimization progress, normalized to 50 iterations/epochs. Overall, medium, decreasing step-sizes performed best in optimization, showing step-sizes similar to the full-batch optimizer Ipopt. Constant learning rates lead to almost constant step-sizes, even for mini-batch algorithms, which are termed adaptive in literature, i.e., RMSProp and Adam. Shading depict percentiles for 20% and 80% of optimizer traces, solid lines depict medians. **A** Step-sizes for Fujita model at medium, decreasing learning rate. **B** Step-sizes for Bachmann model at medium, decreasing learning rate. **C** Step-sizes for Lucarelli model at medium, decreasing learning rate. **D** Step-sizes for Fujita model at low, decreasing learning rate. **E** Step-sizes for Bachmann model at low, decreasing learning rate. **F** Step-sizes for Lucarelli model at low, decreasing learning rate. **G** Step-sizes for Fujita model at constant learning rate. **H** Step-sizes for Bachmann model at constant learning rate. **I** Step-sizes for Lucarelli model at constant learning rate.

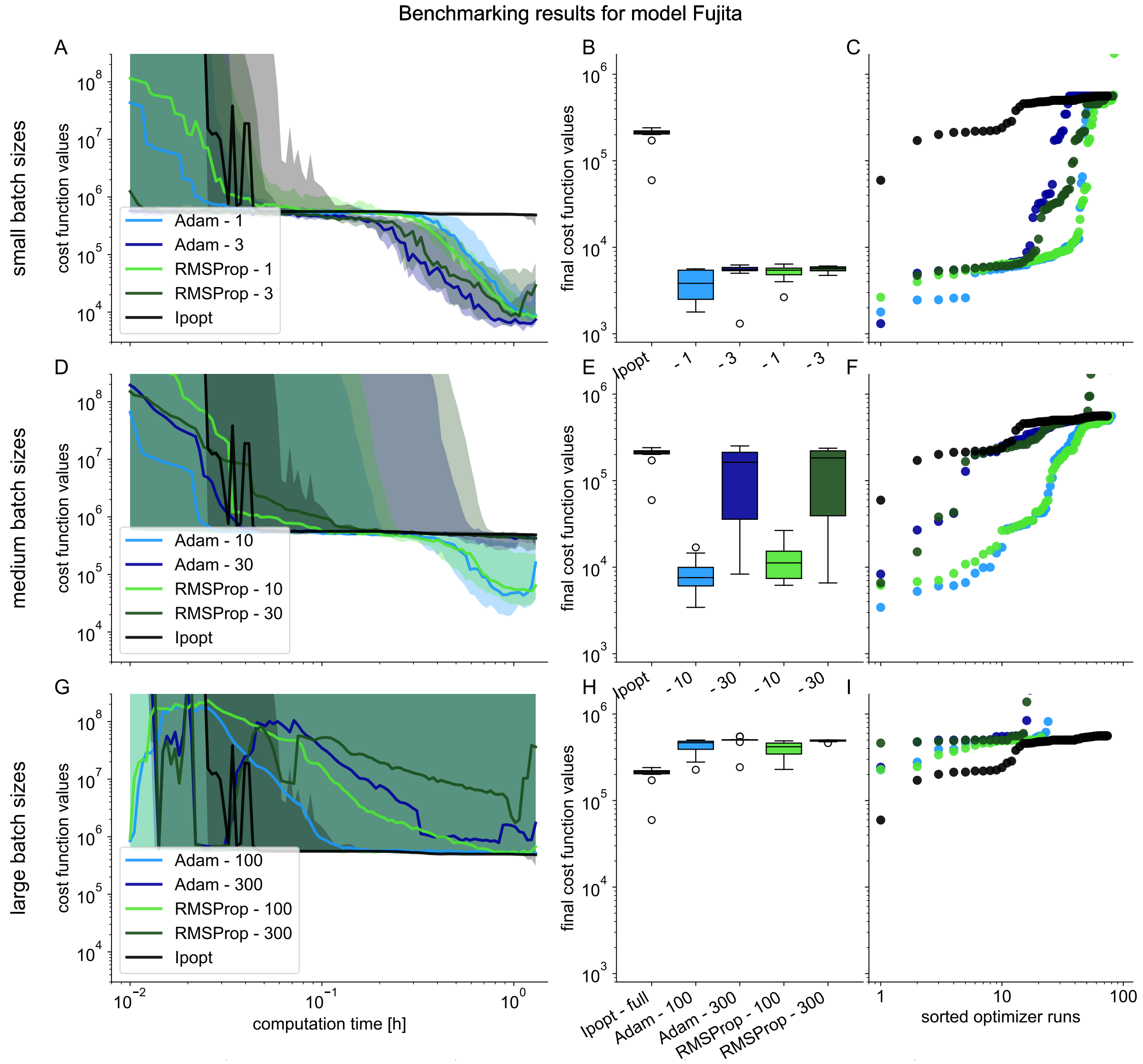


Figure 5: Small mini-batch sizes show best performance for mini-batch optimization on Fujita example model (Supplementary to Fig. 2 of main manuscript). Mini-batch optimizers using different mini-batch sizes were benchmarked against full-batch optimizer Ipopt on Fujita example model. Objective function values along optimization traces were extrapolated from mini-batch to the full dataset, smoothed over multiple optimization steps to filter the stochasticity of the mini-batches, and interpolated for a mesh of computation time points. Shading depicts percentiles for 10% and 40% of optimizer traces, solid lines depict percentiles for 20%. Final objective function values for box and waterfall plots were evaluated at final point of optimization trajectory. **A** Traces of objective function value during optimization for small mini-batch sizes. **B** Box plots of the 10 best final objective function values for small mini-batch sizes. **C** Waterfall plot of all final objective function values for small mini-batch sizes. **D** Traces of objective function value during optimization for medium mini-batch sizes. **E** Box plots of the 10 best final objective function values for medium mini-batch sizes. **F** Waterfall plot of all final objective function values for medium mini-batch sizes. **G** Traces of objective function values during optimization with large mini-batch sizes. **H** Box plots of the 10 best final objective function values with large mini-batch sizes. **I** Waterfall plot of all final objective function values with large mini-batch sizes.

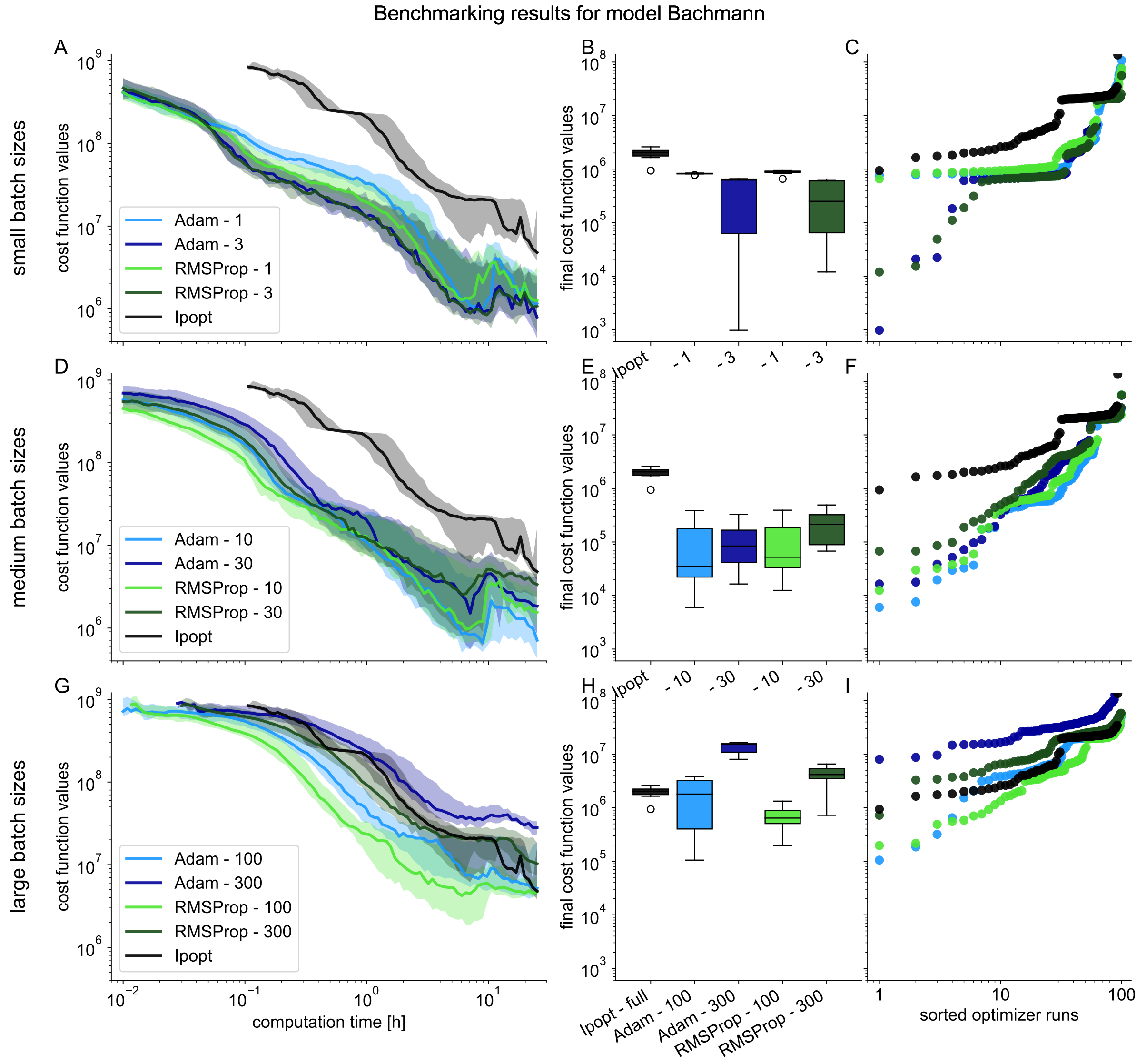


Figure 6: Small mini-batch sizes show best performance for mini-batch optimization on Bachmann example model (Supplementary to Fig. 2 of main manuscript). Mini-batch optimizers using different mini-batch sizes were benchmarked against full-batch optimizer Ipopt on Bachmann example model. Objective function values along optimization traces were extrapolated from mini-batch to the full dataset, smoothed over multiple optimization steps to filter the stochasticity of the mini-batches, and interpolated for a mesh of computation time points. Shading depicts percentiles for 10% and 40% of optimizer traces, solid lines depict percentiles for 20%. Final objective function values for box and waterfall plots were evaluated at final point of optimization trajectory. **A** Traces of objective function value during optimization for small mini-batch sizes. **B** Box plots of the 10 best final objective function values for small mini-batch sizes. **C** Waterfall plot of all final objective function values for small mini-batch sizes. **D** Traces of objective function value during optimization for medium mini-batch sizes. **E** Box plots of the 10 best final objective function values for medium mini-batch sizes. **F** Waterfall plot of all final objective function values for medium mini-batch sizes. **G** Traces of objective function values during optimization with large mini-batch sizes. **H** Box plots of the 10 best final objective function values with large mini-batch sizes. **I** Waterfall plot of all final objective function values with large mini-batch sizes.

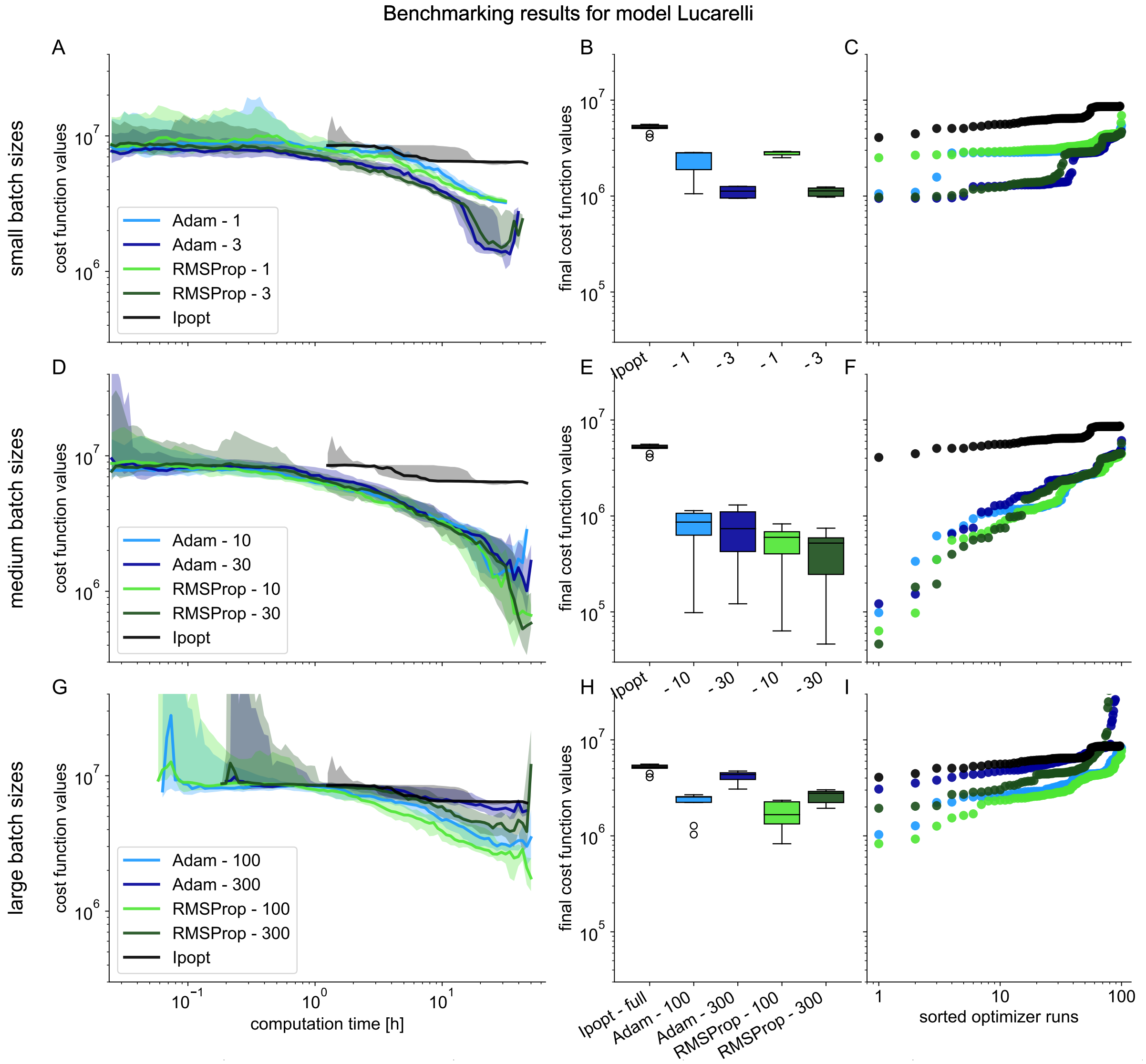


Figure 7: Medium mini-batch sizes show best performance for mini-batch optimization on Lucarelli example model (Supplementary to Fig. 2 of main manuscript). Mini-batch optimizers using different mini-batch sizes were benchmarked against full-batch optimizer Ipopt on Lucarelli example model. Objective function values along optimization traces were extrapolated from mini-batch to the full dataset, smoothed over multiple optimization steps to filter the stochasticity of the mini-batches, and interpolated for a mesh of computation time points. Shading depicts percentiles for 10% and 40% of optimizer traces, solid lines depict percentiles for 20%. Final objective function values for box and waterfall plots were evaluated at final point of optimization trajectory. **A** Traces of objective function value during optimization for small mini-batch sizes. **B** Box plots of the 10 best final objective function values for small mini-batch sizes. **C** Waterfall plot of all final objective function values for small mini-batch sizes. **D** Traces of objective function value during optimization for medium mini-batch sizes. **E** Box plots of the 10 best final objective function values for medium mini-batch sizes. **F** Waterfall plot of all final objective function values for medium mini-batch sizes. **G** Traces of objective function values during optimization with large mini-batch sizes. **H** Box plots of the 10 best final objective function values with large mini-batch sizes. **I** Waterfall plot of all final objective function values with large mini-batch sizes.

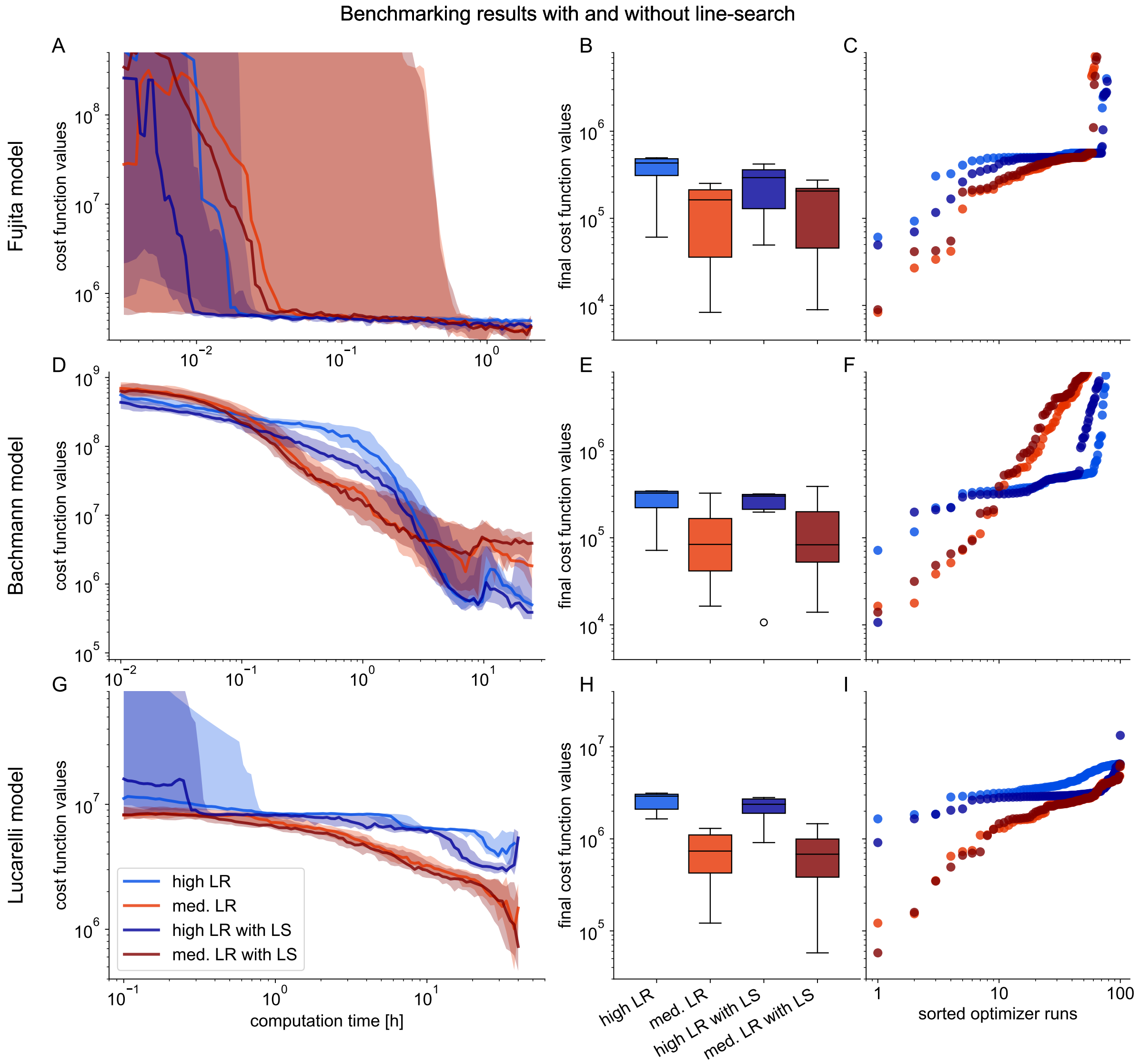


Figure 8: Additional line-search has positive effect for mini-batch optimization methods at high learning rates (Supplementary to Fig. 3 of main manuscript). Optimizations at high and medium learning rates are depicted for small- to medium-scale models. Objective function values along optimization traces were extrapolated from mini-batch to the full dataset, smoothed over multiple optimization steps to filter the stochasticity of the mini-batches, and interpolated for a mesh of computation time points. Shading depicts percentiles for 10% and 40% of optimizer traces, solid lines depict percentiles for 20%. Final objective function values for box and waterfall plots were evaluated at final point of optimization trajectory. **A** Traces of objective function values during optimization for Fujita model. **B** Box plots of the 10 best final objective function values for Fujita model. **C** Waterfall plot of all final objective function values for Fujita model. **D** Traces of objective function values during optimization for Bachmann model. **E** Box plots of the 10 best final objective function values for Bachmann model. **F** Waterfall plot of all final objective function values for Bachmann model. **G** Traces of objective function values during optimization for Lucarelli model. **H** Box plots of the 10 best final objective function values for Lucarelli model. **I** Waterfall plot of all final objective function values for Lucarelli model.

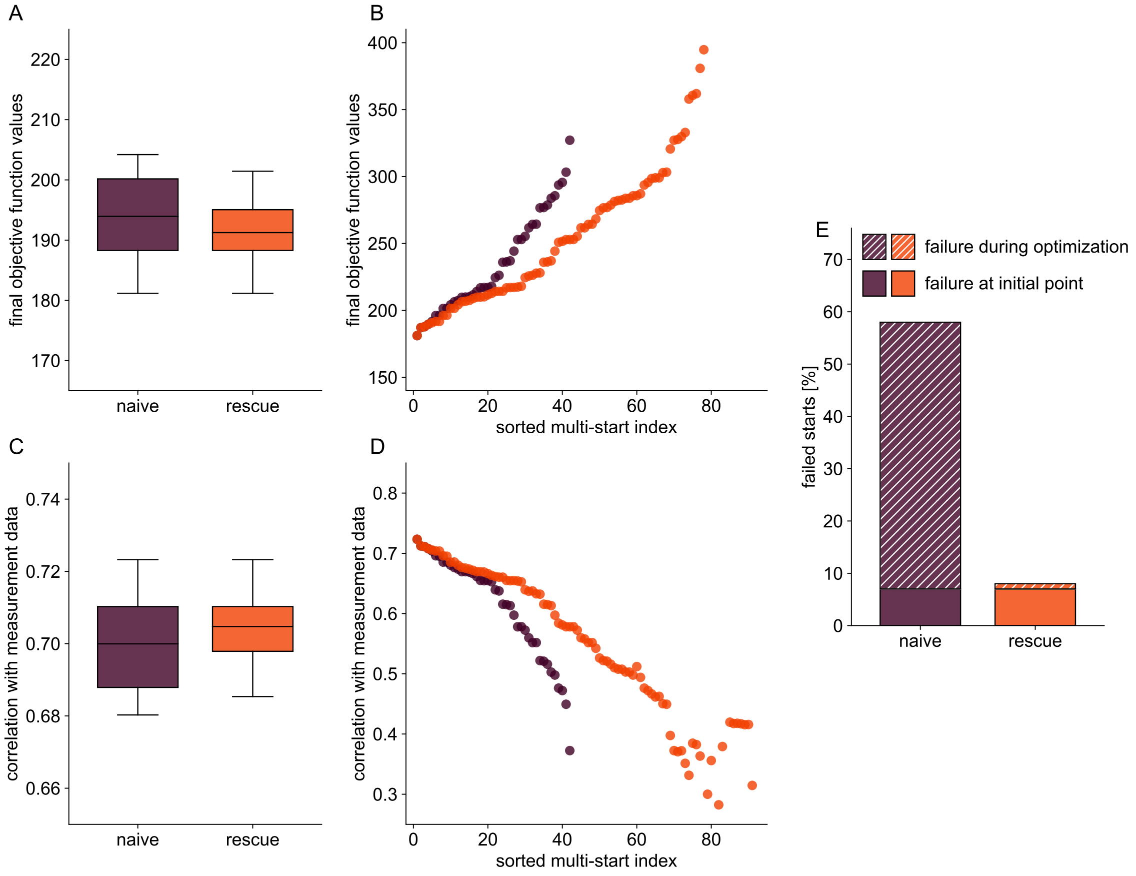


Figure 9: Rescue functionality reduces failure rate for mini-batch optimization on large-scale ODE model (Supplementary to Fig. 4 of main manuscript). Optimization results compared for Adam, mini-batch size 100, at low, decreasing learning rate, with and without rescue functionality. **A** Box plots with final objective function values of the 10 best optimization results. **B** Waterfall plot with final objective function values of optimization results. **C** Box plots with values of correlation between model simulation and measurement data of the 10 best optimization results. **D** Waterfall plot with values of correlation between model simulation and measurement data of optimization results. **E** Bar plot of optimization run failure due to non-integrability of the ODE with and without rescue functionality, indicating failure at initial point of optimization.

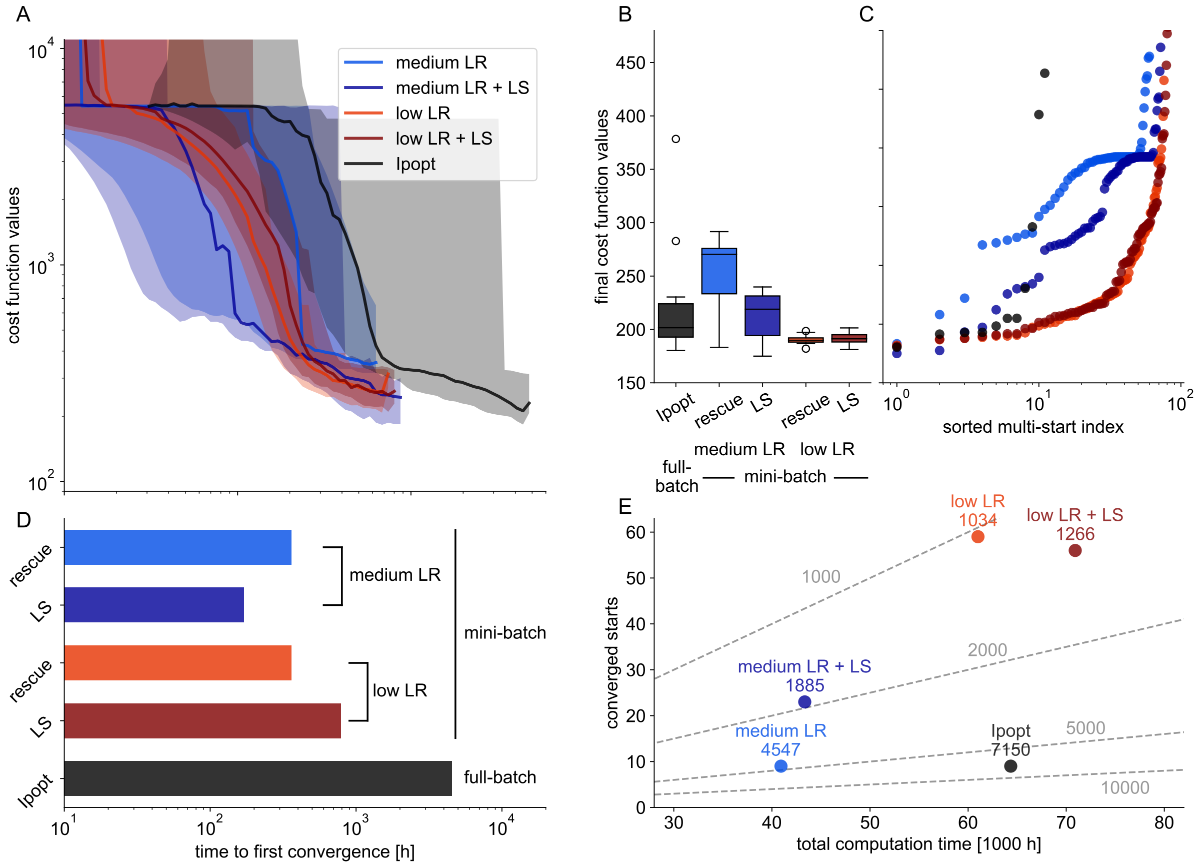


Figure 10: Line-search markedly improves optimization performance for medium learning rates, without substantially obstructing optimization performance for low learning rates on large-scale application example (Supplementary to Fig. 5 of main manuscript). Different metrics of optimization performance are depicted, mini-batch optimization with Adam at mini-batch size 100 was benchmarked for different learning rates with and without additional line-search against full-batch optimizer Ipopt. **A** Traces of objective function values during optimization for the ten best starts plotted against computation time. Objective function values along optimization traces were extrapolated from mini-batch to the full dataset, smoothed over multiple optimization steps to filter the stochasticity of the mini-batches, and interpolated for a mesh of computation time points. Shading depicts the full envelope of the ten best optimizer traces, solid lines depict the medians. **B** Box plots of final objective function values of ten best optimization runs, evaluated at final point of optimization. **C** Waterfall plots of final objective function values of all optimization runs, evaluated at final point of optimization. **D** Bar plots of computation time, until first start converged. **E** Visualization of converged starts against accumulated total computation time showing the computation time per converged start (the lower the value, the better the optimization performance).

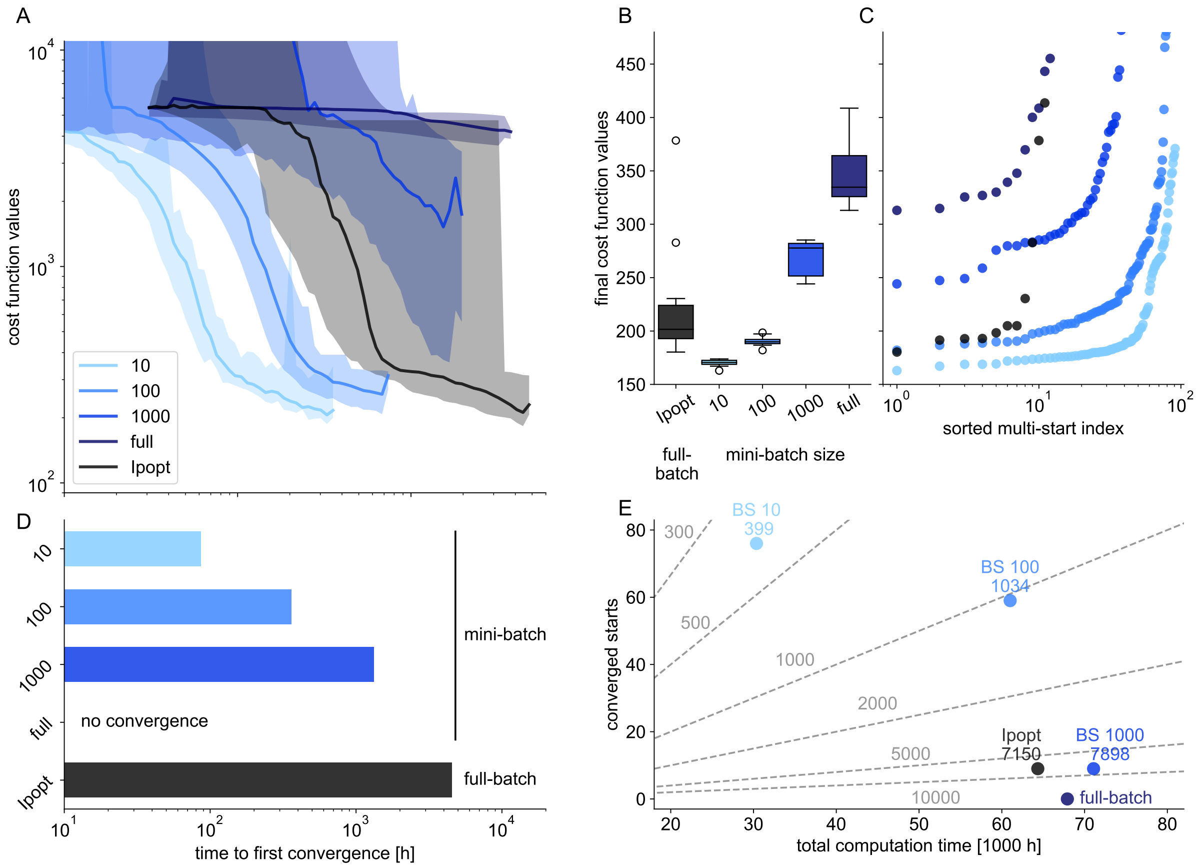


Figure 11: Small mini-batch sizes improve optimization performance for mini-batch optimization on large-scale application example (Supplementary to Fig. 6 of main manuscript). Different metrics of optimization performance are depicted, mini-batch optimization with Adam at mini-batch size 100 was benchmarked for different mini-batch sizes at low learning rates without additional line-search against full-batch optimizer Ipopt. **A** Traces of objective function values during optimization for the ten best starts plotted against computation time. Objective function values along optimization traces were extrapolated from mini-batch to the full dataset, smoothed over multiple optimization steps to filter the stochasticity of the mini-batches, and interpolated for a mesh of computation time points. Shading depicts the full envelope of the ten best optimizer traces, solid lines depict the medians. **B** Box plots of final objective function values of ten best optimization runs, evaluated at final point of optimization. **C** Waterfall plots of final objective function values of all optimization runs, evaluated at final point of optimization. **D** Bar plots of computation time, until first start converged. **E** Visualization of converged starts against accumulated total computation time showing the computation time per converged start (the lower the value, the better the optimization performance).

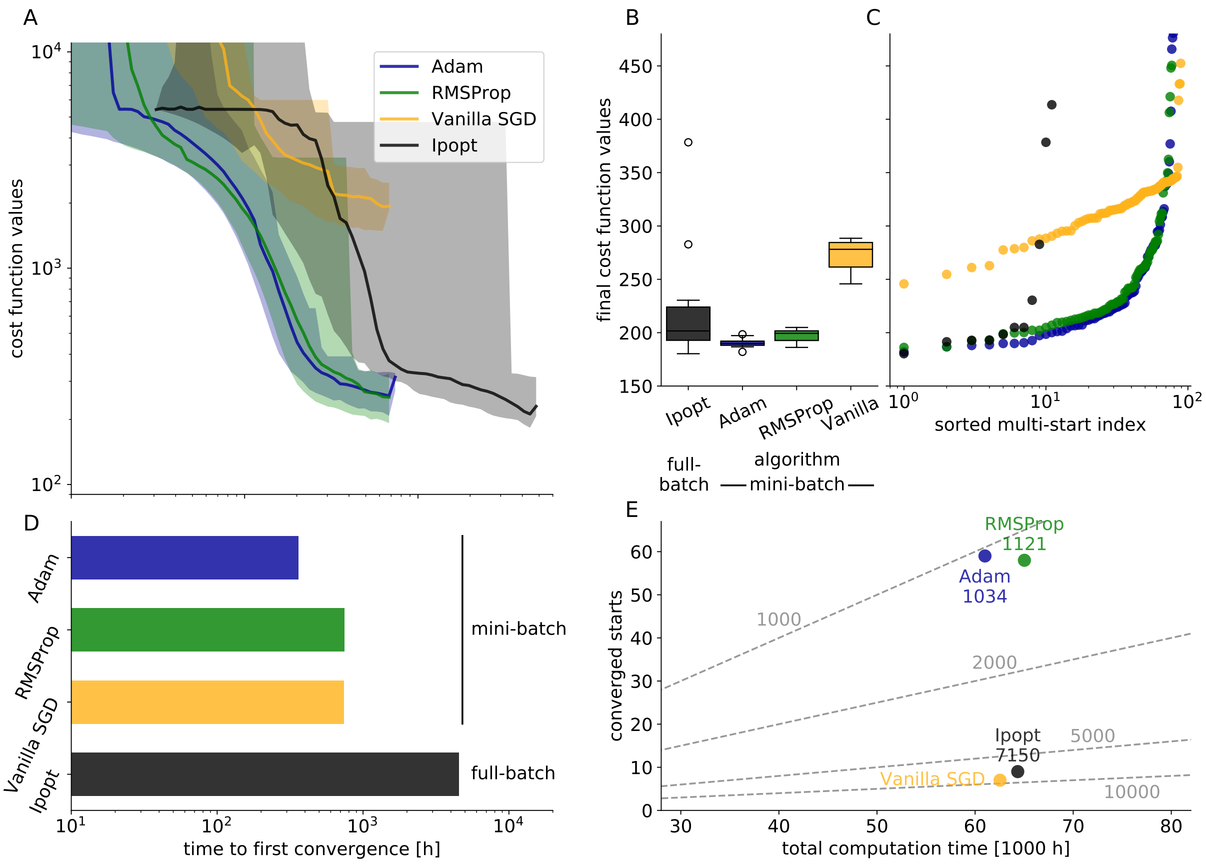


Figure 12: Adam and RMSProp algorithm showing comparable optimization performance on large-scale application example (Supplementary to Fig. 6 of main manuscript). Different metrics of optimization performance are depicted, mini-batch optimization at mini-batch size 100 and was benchmarked for different mini-batch algorithms at low learning rates against full-batch optimizer Ipopt. To ensure similar step-sizes in optimization for all optimizers, we used low learning rates for RMSProp and Adam and a high learning rate ($10^0$ to $10^{-3}$) for Vanilla SGD. **A** Traces of objective function values during optimization for the ten best starts plotted against computation time. Objective function values along optimization traces were extrapolated from mini-batch to the full dataset, smoothed over multiple optimization steps to filter the stochasticity of the mini-batches, and interpolated for a mesh of computation time points. Shading depicts the full envelope of the ten best optimizer traces, solid lines depict the medians. **B** Box plots of final objective function values of ten best optimization runs, evaluated at final point of optimization. **C** Waterfall plots of final objective function values of all optimization runs, evaluated at final point of optimization. **D** Bar plots of computation time, until first start converged. **E** Visualization of converged starts against accumulated total computation time showing the computation time per converged start (the lower the value, the better the optimization performance).
